# Mitochondrial-targeted therapy with elamipretide preserves cardiac function and prevents late mortality in murine polymicrobial sepsis

**DOI:** 10.64898/2026.07.03.736409

**Authors:** Jennie Vu, Cory S. Wagg, Claudia Holody, Anson Wong, Takdeer Baidwan, Maryam Lo, Ibrahim Khodabocus, Si Ning Liu, Kimberly Macala, Antoine Dufour, John Ussher, Helene Lemieux, Gary D. Lopaschuk, Stephane L. Bourque

## Abstract

Sepsis-induced cardiac dysfunction (SICD) occurs in nearly half of septic patients, is associated with increased mortality, and lacks targeted therapy. Emerging evidence implicates impaired mitochondrial function and metabolic inflexibility as central contributors to myocardial depression. Here, we characterized SICD in a murine model of polymicrobial sepsis and evaluated the therapeutic potential of the cardiolipin-stabilizing peptide elamipretide (Ela). Sepsis induced marked impairments in cardiac performance, accompanied by reductions in cardiac cardiolipin content, impaired mitochondrial respiratory capacity localized to complex I, and altered substrate utilization. Integration of stable isotope metabolic flux tracing with lipidomic, metabolomic, and proteomic analyses identified a convergent metabolic bottleneck at the level of the electron transport system. This defect was associated with upstream accumulation of acetyl-CoA, Co-A esters, and ketone bodies, consistent with impaired oxidative flux and energetic failure. Administration of a single early dose of Ela restored cardiolipin content, complex I function, normalized metabolic flux, improved cardiac function during both acute sepsis and recovery, and completely prevented late sepsis-related mortality. These findings identify cardiolipin-dependent mitochondrial dysfunction as a central pathogenic mechanism underlying SICD and position mitochondrial-targeted therapy as a promising therapeutic strategy in sepsis.

**Graphical Abstract:** 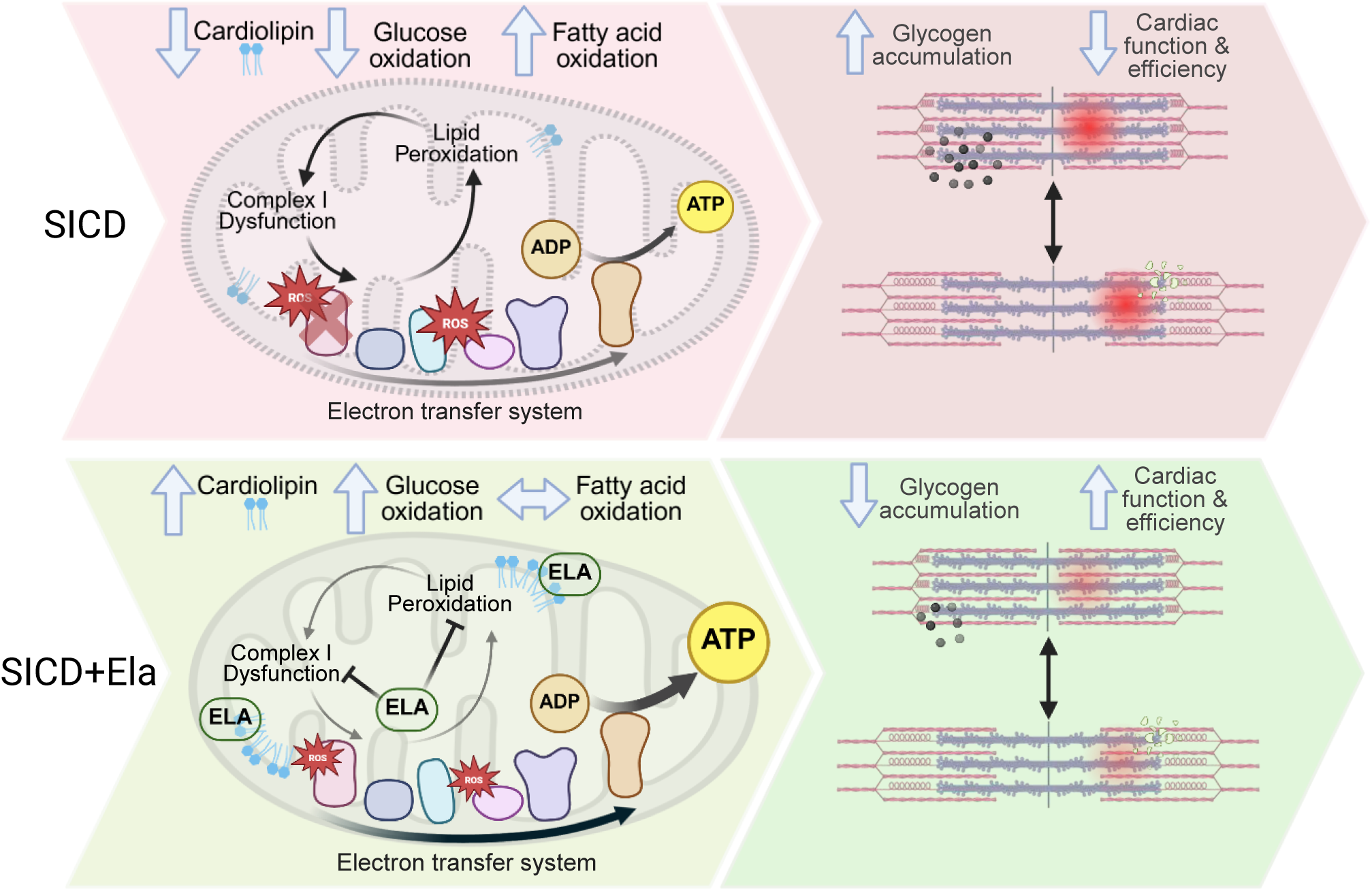

**Ela improves SICD by stabilizing cardiolipin species and improving mitochondrial complex I function.** SICD depicted in red denotes conditions altered compared to healthy control cardiomyocyte, SICD+Ela depicted in green denotes changes relative to SICD. SICD, sepsis-induced cardiac dysfunction; ELA, elamipretide; ADP, adenosine diphosphate; ATP, adenosine triphosphate; ROS, reactive oxygen species.

## Introduction

Sepsis is a dysregulated host response to infection that leads to life-threatening organ dysfunction.^1^ It is a leading cause of intensive care unit admission and mortality^2,3^ and estimates from the Global Burden of Disease Study suggest that sepsis contributes to approximately one in five deaths worldwide.^4^ Among affected organs, the heart is particularly vulnerable. Sepsis-induced cardiac dysfunction (SICD), characterized by reduced biventricular contractility and impaired left ventricular (LV) relaxation, occurs in nearly half of septic patients and is associated with increased mortality^5,6^ and persistent adverse health outcomes among survivors.^7,8^ Despite its clinical importance, the mechanisms underlying SICD remain incompletely understood.

Several observations suggest that myocardial bioenergetic failure plays a central role in SICD. Clinical studies have reported minimal cardiomyocyte death in septic patients with cardiac dysfunction^9^ as well as preserved coronary blood flow and adequate, or even elevated, myocardial oxygen delivery.^10–12^ These findings indicate that impaired oxygen utilization, rather than oxygen supply, may underlie septic myocardial dysfunction. Consistent with this hypothesis, experimental models of sepsis demonstrate impaired mitochondrial oxidative phosphorylation (OXPHOS), reduced respiratory complex activity, increased reactive oxygen species production, and structural mitochondrial abnormalities, including cristae disruption and swelling.^9,13–15^ In parallel, endotoxemia and sepsis models exhibit alterations in cardiac substrate metabolism, including reduced myocardial glucose oxidation^16^ and altered fatty acid oxidation^10,17^—the heart’s primary source of ATP generation.^18^ Collectively, these findings implicate mitochondrial dysfunction as a central contributor to SICD. However, the upstream mechanisms linking sepsis to impaired OXPHOS, altered substrate utilization and contractile dysfunction remain poorly defined.

Cardiolipin (CL) is a mitochondria-specific tetra-acyl phospholipid that comprises approximately 20% of the phospholipid content of the inner mitochondrial membrane.^19^ Owing to its unique conical structure, cardiolipin plays a critical role in maintaining membrane curvature and cristae architecture, stabilizing respiratory chain complexes, and supporting efficient OXPHOS. Disruption of cardiolipin content or remodeling impairs mitochondrial function in disorders ranging from Barth syndrome^20^ to chronic heart failure^21–25^ Cardiolipin is also particularly susceptible to oxidative damage because of its high content of polyunsaturated fatty acyl chains and its proximity to the electron transport system. Recent studies demonstrating altered cardiolipin remodeling in lipopolysaccharide-exposed cardiomyocytes,^26^ suggest that cardiolipin dysregulation may mechanistically link septic stress and mitochondrial dysfunction in the heart.

Because metabolic remodeling likely precedes irreversible structural injury in the septic heart, interventions targeting mitochondrial bioenergetics may represent a promising therapeutic strategy. Elamipretide (Ela; also known as SS-31 or MTP-131) is a mitochondria-targeted cationic-aromatic tetrapeptide that selectively binds cardiolipin and stabilizes mitochondrial membrane structure and function.^27^ Elamipretide has recently received regulatory approval for the treatment of Barth syndrome and has demonstrated beneficial effects in experimental models of heart failure, ischemia–reperfusion injury, and age-related cardiac dysfunction.^28–31^ However, its potential therapeutic role in SICD has not been evaluated.

Based on these observations, we hypothesized that sepsis disrupts cardiolipin-dependent mitochondrial function in the heart, resulting in impaired electron transport, altered substrate utilization, and myocardial energetic failure that contributes directly to contractile dysfunction. To test this hypothesis, we used a murine model of polymicrobial sepsis to define the relationship among cardiolipin remodeling, mitochondrial respiratory dysfunction, and cardiac reprogramming. We further examined whether treatment with elamipretide could restore mitochondrial function, improve cardiac performance, and modify both acute and post-recovery outcomes. Our findings identify cardiolipin-dependent mitochondrial dysfunction as a key mechanism underlying SICD and provide strong evidence supporting mitochondrial-targeted therapy as a potential treatment strategy for sepsis.

## Methods and Materials

### Polymicrobial peritonitis mouse model of sepsis

Ten to twelve-week-old male mice (C57BL/6N, Charles River, Kingston, NY, USA) were injected intraperitoneally (i.p.) with either cecal slurry (CS; 0.55 mg/kg) or vehicle (5% dextrose,10% glycerol) (Fig. 1A). CS was prepared from adult male Sprague Dawley rats as previously described,^32^ aliquoted and stored at −80℃ for up to 1 year; each aliquot underwent a single freeze-thaw cycle before use. At 1h post-inoculation (T1h) mice received i.p. Ela (10 mg/kg) or PBS vehicle. Pain management was provided at 4h (T4h) via subcutaneous (s.c.) buprenorphine (0.5 mg/kg). Mice were assessed hourly using the murine sepsis score (MSS) until T12h, and every 12h thereafter until 72h (T72h). MSS evaluates appearance, activity, response to stimulus, and respiration.^32^

**Figure 1.**
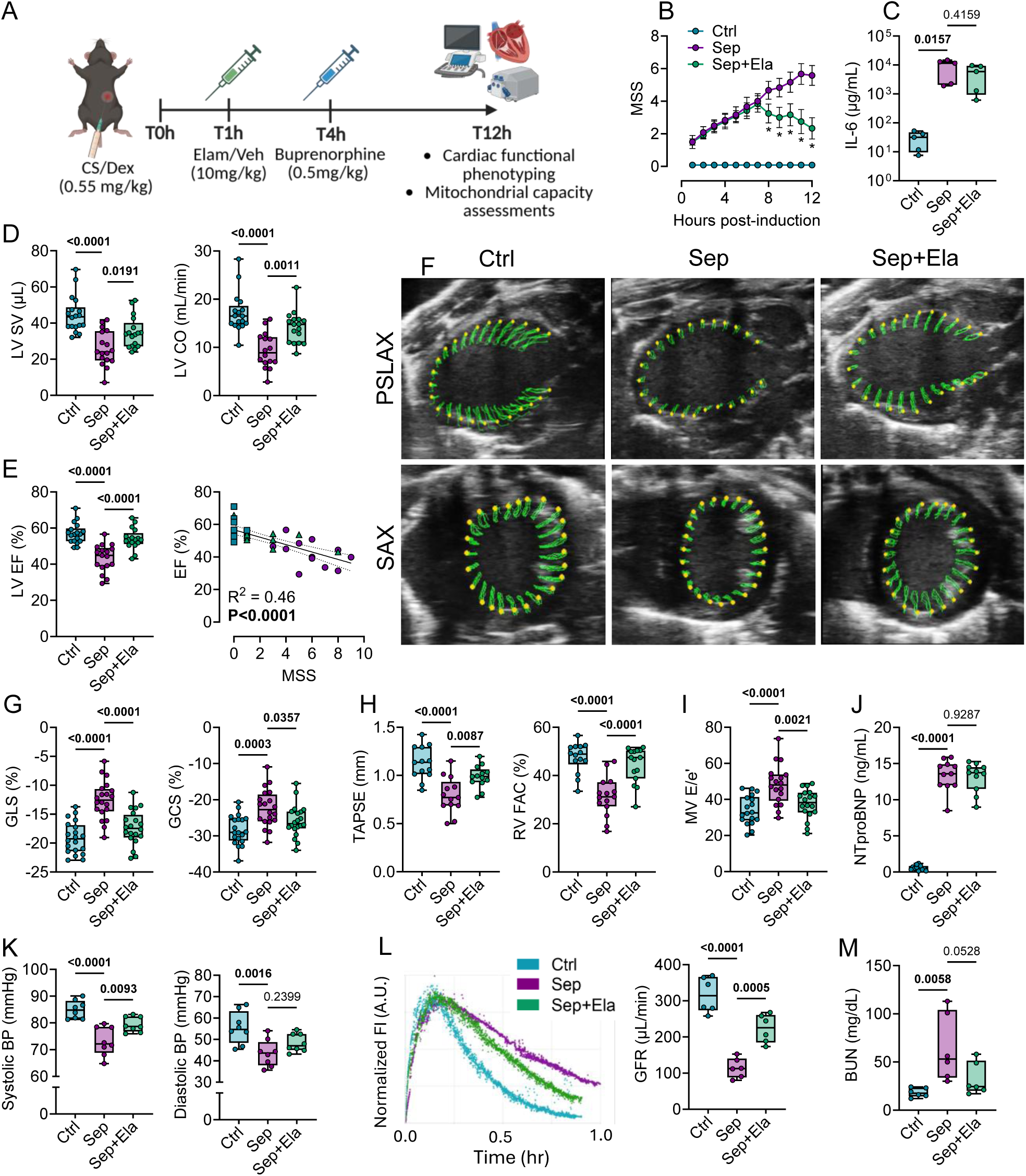
Elamipretide attenuates cardiac, hemodynamic, and renal dysfunction in a murine cecal slurry model of sepsis. **A**. Experimental timeline. Adult mice were inoculated with cecal slurry to induce peritonitis at time 0 (T0h), followed by intraperitoneal administration of elamipretide (Sep+Ela) or vehicle (Sep) at 1 hour (T1h). Cardiac and hemodynamic function were assessed by echocardiography and carotid artery catheterization, and renal function by transdermal GFR measurement, at 12 hours post-induction. **B**. Murine sepsis score (MSS) assessed at serial timepoints across 12 hours, incorporating appearance, orbital tightening, activity, response to stimulus, and breathing quality. **C**. Plasma interleukin-6 (IL-6) concentrations at T12h (log-transformed). **D**. Left ventricular stroke volume (SV) and cardiac output (CO) assessed by echocardiography. **E**. Left ventricular ejection fraction (EF; left) and correlation of LV EF with MSS at T12h (right). **F**. Representative speckle-tracking echocardiographic images depicting LV wall motion used to quantify global longitudinal strain (GLS) in the parasternal long-axis (PSLAX) view (top) and global circumferential strain (GCS) in the parasternal short-axis (SAX) view (bottom). Green overlays trace endocardial motion across two to three cardiac cycles. **G**. Quantification of LV GLS and GCS. **H**. Tricuspid annular plane systolic excursion (TAPSE) and right ventricular fractional area change (RV FAC). **I**. Mitral valve E/e’ ratio as an index of left ventricular filling pressure. J. Plasma NT-proBNP concentrations. **K**. Systolic and diastolic blood pressure measured by carotid artery catheterization using a pressure micromanometer. **L**. Representative FITC-sinistrin elimination curves measured by transcutaneous fluorescence detection (left) and corresponding glomerular filtration rate (GFR) calculated from the body weight-adjusted elimination half-life (right). **M**. Blood urea nitrogen (BUN) concentrations at T12h (log-transformed). Data are presented as box plots with individual data points overlaid; boxes represent interquartile ranges (IQR, 25^th^-75^th^ percentile), horizontal lines indicate median, maximum and minimum values (n = 5-20 per group). All comparisons were performed by one-way ANOVA with Sidak’s multiple comparison test; exact p values are indicated between groups. Ctrl, naive healthy control mice; Sep, cecal slurry-induced septic mice administered vehicle; Sep+Ela, cecal slurry-induced septic mice administered elamipretide.

Separate cohorts of mice underwent functional assessments at T12h and 10 days (T10d), including echocardiography, pressure-volume loop analysis, and glomerular filtration rate assessments. Mice in the T10d cohorts received s.c. antibiotics (imipenem-cilastatin, 25 mg/kg) and Ringer’s lactate (s.c., 15 mL/kg) at T12h and every 12 hours thereafter until T72h. Following functional assessments, mice were euthanized by exsanguinated via carotid artery transection. Blood was collected in heparinized tubes, and excised hearts were cardiopleged in 0.1M KCl prior to separation of the left ventricle (LV) and right ventricle (RV) for either transmission electron microscopy (TEM) fixation, or flash-freezing in liquid nitrogen.

### Echocardiography

Electrocardiography-gated, speckle-tracking, transthoracic echocardiography was performed at T12h or T10d using a MX550S 25-55MHz transducer probe (Vevo 3100, VisualSonics) by an investigator blinded to group assignment. Parasternal long- and short-axis B- and M-mode recordings were acquired. Subcostal apical 4-chamber views enabled pulsed-wave and tissue Doppler imaging of the mitral valve (MV) and tricuspid valve (TV), tricuspid annular plane systolic excursion (TAPSE), and RV fractional area change (RV FAC). Data analysis was conducted using Vevo Lab 5.8.0. (Fujifilm, VisualSonics). Global longitudinal strain (GLS) and global circumferential strain (GCS) were determined using the VevoStrain function, averaging values from four cardiac cycles across two images for each parameter.

### Pressure volume (PV) loop assessments and blood analysis

PV loop assessments were performed at T12h and T10d using the MPVS Ultra system and PowerLab 8/35 (Millar, ADInstruments). Mice were anesthetized with 1.5-2% isoflurane in 100% oxygen, and the right carotid artery was catheterized with a 1.0F PV catheter (PVR-1045). Baseline arterial pressure was recorded for 5min after a 10min equilibration period. The catheter was then advanced into the LV cavity to record simultaneous pressure and volume for 5min. Hemodynamic challenges included inferior vena cava occlusion (5s) to reduce preload and aortic occlusion (5s) to increase afterload after 30s stabilization. An investigator blinded to group assignment used LabChart 8.1.28 to analyze 100 cardiac cycles and responses to hemodynamic challenges.

### Glomerular filtration rate (GFR)

Mice were anesthetized (1.5-2% isoflurane in 100% inspired oxygen) and dorsally depilated. A battery-powered preclinical transcutaneous fluorescence monitor (Model MX2904, MediBeacon, Germany) was attached to the skin. FITC-sinistrin (0.06mg/g BW, retro-orbital) was administered, and mice recovered from anesthesia. Data were collected for ≥1h and analyzed with MediBeacon Lab 3 and Studio 3 software. Plasma curves were fitted to a 3-compartment model with baseline correction; GFR was calculated from FITC-sinistrin half-life using an empirically derived formula for mice.^33^

### High resolution mitochondrial respirometry

Fresh LV lateral wall tissue (∼5mg in 2mL) was homogenized in Mir05 for assessment of mitochondrial respiratory capacity using an Oxygraph-2K (Oroboros Instruments) as described.^34^ Two substrate-uncoupler-inhibitor-titration (SUIT) protocols were performed, initiated with either pyruvate (5mM) and malate (2mM) or with palmitoylcarnitine (10μM) and malate (0.1mM). Both protocols then received sequential additions of ADP (2.5mM), cytochrome *c* (10μM), glutamate (10mM), succinate (10mM), rotenone (1μM), antimycin A (5μM), ascorbate (2mM) and tetramethylphenylenediamine (0.5mM), and sodium azide (100mM) for chemical background correction. Outer mitochondrial membrane integrity was assessed by cytochrome *c* control efficiency, calculated as the fractional increase in O_2_ flux following cytochrome *c* addition relative to the flux measured after cytochrome *c* addition and expressed as a percentage.

### Isolated working heart perfusion

A subset of mice was anesthetized with sodium pentobarbital (i.p., 1 g/kg) at T12h. Hearts were excised and perfused in a working heart apparatus under a left atrial preload of 11.5 mmHg and an aortic afterload of 50 mmHg in Krebs-Henseleit solution supplemented with 0.8mM palmitate bound to 3% albumin, 5mM glucose, and 0.6mM β-hydroxybutyrate (βOHB).^35^ Radiolabeled substrates [9,10-³H]-palmitate and [U-¹⁴C]-glucose were used to measure fatty acid and glucose oxidation, respectively. Glucose oxidation rates were assessed by measuring ^14^CO_2_ production, and palmitate oxidation was assessed by measuring ^3^H_2_O production, following established protocols. A second cohort used [5-^3^H]-glucose and [3-^14^C]-βOHB to assess glycolytic flux and ketone oxidation by quantitative extraction of ^3^H_2_O and ^14^CO_2_ respectively.^36^

### Proteomics, metabolomics, and lipidomics

LV middle-anterolateral tissue (20mg) was snap-frozen in liquid nitrogen and analyzed by data-independent acquisition quantitative shotgun proteomics.^37^ Spectra were aligned to the mouse UniProt database. Functional analyses were performed using Metascape and STRING v.11.5 for pathway enrichment (FDR <0.05). Protein abundance changes are presented as Log2 fold changes. For metabolomics, tissue homogenates (3 volumes extraction buffer) were centrifuged, and supernatants analyzed using targeted LC-MS/MS (AB Sciex 5500 QTrap). Up to 268 metabolites were quantified using multiple reaction monitoring with isotope-labeled internal standards. Organic acids were derived from 3-nitrophenylhydrazine prior to analysis. For lipidomic profiling, lipids were extracted using a modified Folch protocol, spiked with internal standards, and analyzed via Agilent 1290 LC coupled to a 6546 QTOF-MS using a CSH C18 column. Data acquisition included MS/MS for identification, with quality control injections interspersed. Features detected in ≥80% of samples were retained.

### Transmission electron microscopy

LV middle inferolateral wall tissue was cut into 1mm^3^ cubes, fixed in cold 2% paraformaldehyde/2.5% glutaraldehyde in 0.1M cacodylate buffer, and imaged by TEM (70-100 images/sample) by a blinded operator.

### Enzyme activity assays

Complex I activity was measured as described.^38^ Frozen LV tissue (3.3mg/mL) was homogenized in imidazole-HCl buffer (50mM, pH 7.5). NADH:ubiquinone oxidoreductase activity was measured spectrophotometrically at 340nm (ε = 6.81mM·cm⁻¹), with rotenone-sensitive activity used to calculate complex I activity. Pyruvate dehydrogenase (PDH) activity was measured as described.^39^ Homogenates were activated with MgCl₂ and mixed with reaction buffer containing NAD^+^, CoA, INT, TPP and lipoamide dehydrogenase. PDH activity was calculated from INT reduction rate (ε = 15.4 mM⁻¹·cm⁻¹). LV ATP was measured using the Abcam ATP Assay Kit (ab83355). Samples were homogenized in 2M perchloric acid, neutralized, and assayed at 570nm.

### Plasma cytokines

Plasma cytokine, chemokine, and growth factor levels were measured using Luminex xMAP multiplexing (Mouse Cytokine/Chemokine 32-Plex, MilliporeSigma) by Eve Technologies (Calgary, Canada). Sensitivities ranged from 0.4 to 10.9pg/mL.

### Western blot

LV tissue lysates were prepared in CelLytic reagent (Millipore-Sigma), resolved on 10-12% SDS-PAGE, transferred to PVDF membranes, and probed for malondialdehyde (Abcam, ab27642), O-GlcNAc (#9875, Cell Signaling Technology, Boston, MA), PDK4 (sc-51806, Santa Cruz), p-PDH (Ser293; #31866. Cell Signaling Technology), PDH (#3205, Cell Signaling Technology), AMPK (#2532, Cell Signaling Technology), p-AMPK (Thr172; #2531, Cell Signaling Technology), and FOXO1 (#2880, Cell Signaling Technology) with nuclear fractionation using the Abcam Nuclear Extraction Kit (ab219177); PARP1 as nuclear-soluble marker, GAPDH as cytoplasmic marker. HRP-conjugated secondary antibodies for chemiluminescent imaging were used for quantification (Image Studio Lite).

### Quantitative Reverse Transcription Polymerase Chain Reaction (RT qPCR)

Total RNA was isolated using TRIzol, chloroform, and isopropanol. cDNA synthesis used 2µg RNA and All-In-One 5X RT MasterMix (abm). qPCR was performed with PowerUp SYBR Green Master Mix on a LightCycler 480 (Roche). Data were analyzed using the 2^−ΔΔCt^ method, normalized to Actb. Primer specificity was verified by melt curves. Primer sequences for all transcripts assessed are provided in **Supplementary Table S1.**

### Statistical analysis

Parametric data across the three groups were analyzed by one-way ANOVA with Šídák’s post hoc multiple-comparisons test, using pre-specified contrasts to evaluate the effects of sepsis (Sep vs Ctrl) and the recovery profile following Ela treatment (Sep+Ela vs Sep). Non-parametric data were analyzed using Kruskal-Wallis with Dunn’s multiple-comparison test. Repeated or time-course assessments were analyzed by 2-way ANOVA with Šídák’s post hoc test. Survival data were analyzed by the log-rank (Mantel-Cox) test. Normality was assessed by the Shapiro-Wilk test, and homogeneity of variance by Bartlett’s test; data that violated these assumptions were log-transformed prior to analysis. Sample sizes were determined a priori (α=0.05, power [1-β]=0.80) for the primary functional endpoint (mitochondrial respiratory capacity) based on variance from our previously published studies.^34^

All omics analyses were conducted under a common confirmatory–exploratory framework applied consistently across proteomics, lipidomics, and targeted central carbon metabolomics. For each assay, one or more primary endpoints were pre-specified prior to analysis, selected on mechanistic grounds from the central hypothesis that sepsis-induced complex I dysfunction restricts mitochondrial substrate oxidation. Primary endpoints were tested directly and are reported with effect sizes (log₂ fold change) and nominal P values, without multiplicity correction, consistent with their status as a small set of pre-specified hypotheses rather than unbiased discovery comparisons. The primary endpoints were: PDK4 (proteomics), the regulatory kinase predicted to suppress PDH activity; cardiolipin species (lipidomics), the inner-membrane phospholipid central to cristae organization and electron transport chain function; and the PDH-axis metabolites pyruvate, acetyl-CoA, and glucose 6-phosphate (metabolomics), reflecting the predicted pyruvate-oxidation bottleneck and upstream glycolytic congestion. All remaining features within each assay were treated as exploratory and are reported with effect sizes and Benjamini–Hochberg FDR-adjusted P values. Two contrasts were evaluated in every assay: Sep vs Ctrl (disease effect) and Sep+Ela versus Sep (rescue effect), with FDR correction applied within each assay and contrast. Analyses were performed in Python 3.14 (SciPy, statsmodels, scikit-learn) and Graphpad Prism v11.02. For all omics analyses, outliers were removed per group per feature using the ROUT method (Q = 1%)^40^; at the sample sizes employed, no features met outlier criteria. Data outputs were log2-transformed and normalized, and differential abundance was assessed by Welch’s t test with Benjamini–Hochberg FDR correction; exploratory significance was defined at FDR<0.01 for proteomics and metabolomics, and FDR<0.05 for lipidomics. Principal component analysis was performed on autoscaled log₂ data with group separation assessed by PERMANOVA (Euclidean distance, 9,999 permutations).

## RESULTS

### Elamipretide mitigates sepsis-induced cardiac dysfunction

Peritoneal injection of cecal slurry (CS) induced illness by T1h, as reflected by increased murine sepsis scores (MSS), that progressed over the following 12h (**Fig. 1A-B**). Increases in the proinflammatory cytokine IL-6 were evident by T4h, the earliest timepoint assessed, and persisted at T24h, with robust systemic inflammation evident by T12h (**Fig. 1C, S1A-B**).

Echocardiography at T12h revealed marked biventricular systolic dysfunction in Sep mice. Compared with vehicle-treated controls, Sep mice exhibited reduced LV stroke volume (SV) and cardiac output (CO), along with reduced ejection fraction (EF) (**Fig. 1D-E**). EF and CO were inversely correlated with MSS (**Fig. 1G, S1C**), indicating that cardiac dysfunction tracked with overall illness severity. Hypokinetic LV wall motion in Sep mice corresponded to reduced global longitudinal strain (GLS) and global circumferential strain (GCS), which also correlated with MSS (**Fig. 1F-G, S1D**). RV systolic function was similarly impaired, as reflected by reduced TAPSE and RV FAC (**Fig. 1H**). Sep mice also exhibited evidence of LV diastolic dysfunction, with an elevated MV E/e’ ratio despite an unchanged MV E/A ratio (**Fig. 1I, S1E**). In contrast, indices of RV diastolic function (tricuspid valve E/A and E/e’) and pulmonary vascular resistance (PAAT/PAET) were unchanged relative to vehicle-treated controls (**Fig. S1F-G**).

Administration of Ela at T1h attenuated illness severity beginning at T8h, but had minimal impact on circulating inflammatory mediators, including IL-6, MCP-1, TNF-α and IL-10 at T12h (**Fig. 1B-C, S1B**). In contrast, Ela markedly improved cardiac performance: Ela-treated Sep mice were protected from declines in LV systolic function, had improved LV wall motion, and demonstrated improved RV systolic function (**Fig. 1D-H**). Ela also normalized LV diastolic function, reducing MV E/e’ (**Fig. 1I**). Despite these improvements, NTproBNP remained elevated in both Sep groups, indicating persistent cardiac stress (**Fig. 1J**). Heart rate and heart weights were unaffected by sepsis or by Ela treatment (**Fig. S1H-I**).

Consistent with septic circulatory dysfunction, Sep mice developed hypotension at T12h, and Ela selectively improved systolic but not diastolic blood pressure (**Fig. 1K**), suggesting that improved cardiac performance contributed more prominently to the hemodynamic response than changes in vascular tone. Sep mice also exhibited renal dysfunction, characterized by reduced GFR, elevated blood urea nitrogen levels (BUN), and increased urinary NGAL (**Fig. 1L-M, S1J**). Ela partially restored GFR and tended to reduce BUN but did not alter urinary NGAL (**Fig. 1L-M, Fig. S1J**), consistent with improved renal hemodynamics rather than direct protection against intrarenal injury.^41^ Finally, Sep mice had elevated lung wet-to-dry weight ratios and hemoconcentration, consistent with pulmonary edema and fluid third-spacing, which were improved with Ela treatment (**Fig. S1K**).

Collectively, these findings demonstrate that CS-induced sepsis rapidly produces severe cardiac and circulatory dysfunction, and that early administration of Ela mitigates these cardiovascular impairments despite persistent systemic inflammation.

### Elamipretide attenuates sepsis-induced impairment of intrinsic cardiac contractile capacity

Because septic mice exhibited hemodynamic changes, reductions in cardiac performance could reflect altered loading conditions rather than intrinsic myocardial dysfunction. We therefore performed pressure–volume (PV) loop analyses at T12h to directly assess load-independent myocardial contractile capacity and determine whether Ela preserves these outcomes (**Fig. 2A**). Sep mice exhibited markedly reduced stroke work (SW), indicating reduced myocardial energy expenditure for blood ejection (**Fig. 2B**). Potential energy was unchanged between groups, resulting in Sep mice having reduced cardiac efficiency, defined as the ratio of SW to total PV area (**Fig. S2A, 2C**). Consistent with impaired intrinsic contractility, the load-independent index of contractility, end-systolic elastance (Ees), was reduced in Sep mice relative to controls (**Fig. 2D**). Arterial elastance (Ea), an indicator of ventricular afterload, was unchanged, indicating that the increased Ea/Ees ratio (ventriculo-arterial coupling; VAC) observed in Sep mice primarily reflected impaired myocardial contractility rather than altered afterload (**Fig. S2B, 2E)**. Load-independent indices derived from preload manipulation further support the presence of impaired contractile capacity. Inferior vena cava occlusion revealed reductions in both preload-recruitable stroke work (PRSW) and preload-recruitable cardiac work (PRCW), the slopes of the relationships between SW or total PV area and end-diastolic volume (EDV), respectively, in Sep mice, confirming diminished intrinsic myocardial contractility independent of loading conditions (**Fig. 2F-G**).

**Figure 2.**
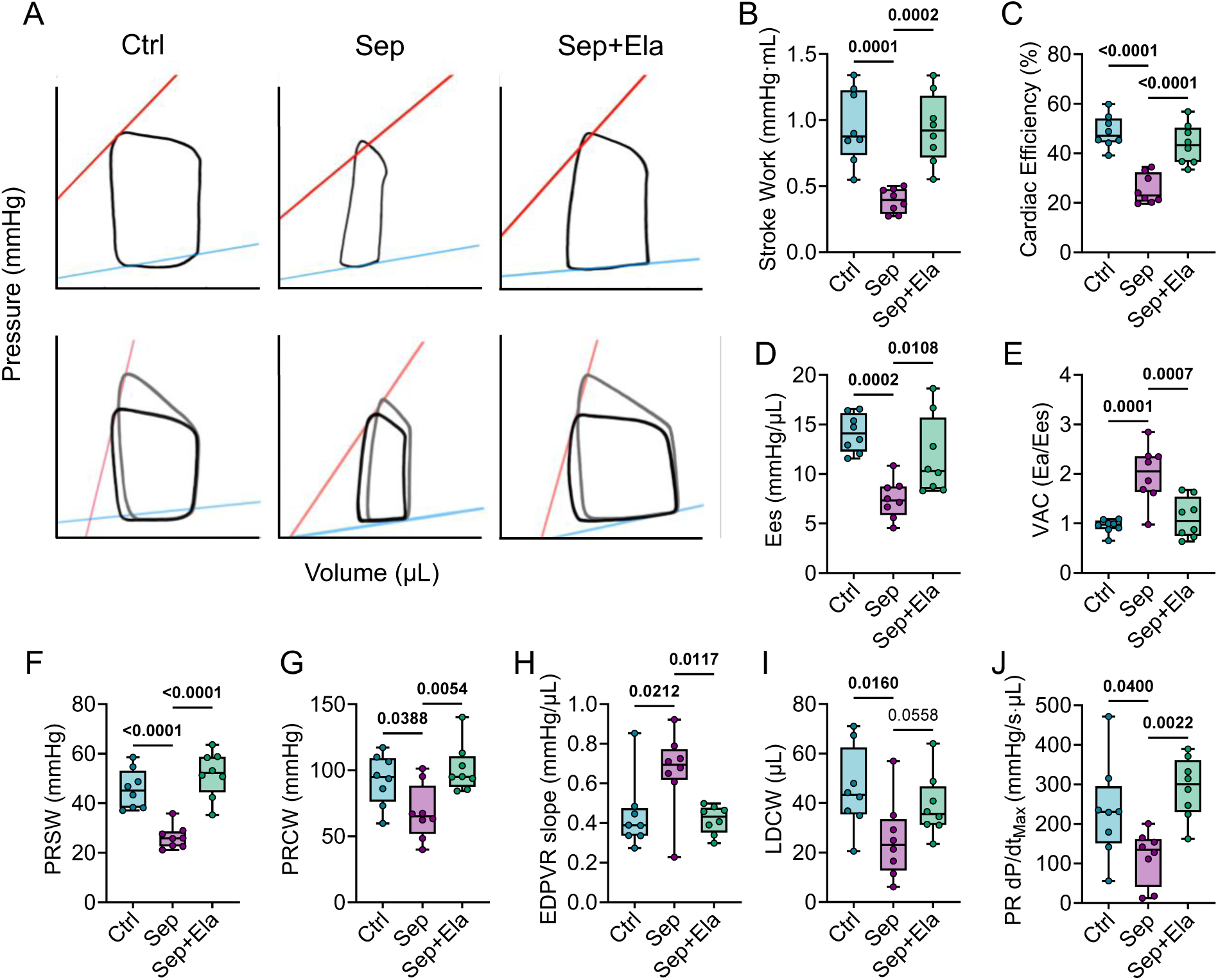
Pressure-Volume loop analysis reveals sepsis-induced impairment of intrinsic cardiac contractile and relaxation function that is partially restored by elamipretide. **A**. Representative left ventricular pressure-volume (PV) loops recorded at baseline (top row) and during aortic occlusion to increase afterload (bottom row) in Control, Septic, and Sep+Ela mice (left to right). The end-systolic pressure-volume relationship (ESPVR; red line) and end-diastolic pressure-volume relationship (EDPVR; blue line) are indicated. **B**. Stroke work (SW), quantified as the area enclosed by the pressure-volume loop. **C**. Cardiac efficiency, defined as the ratio of stroke work to total pressure-volume area (PVA); total PVA comprises stroke work plus potential energy, where potential energy is defined as the region bounded by the ESPVR, EDPVR, and the isovolumetric relaxation line. **D**. End-systolic elastance (Ees), the slope of the ESPVR, as a load-independent index of myocardial contractility. **E**. Ventriculo-arterial coupling (VAC), defined as the ratio of Ea (arterial elastance) to Ees. **F**. Preload-recruitable stroke work (PRSW), defined as the slope of the linear relationship between stroke work and end-diastolic volume (EDV). **G**. Preload-recruitable cardiac work (PRCW), defined as the slope of the relationship between total PVA and EDV. **H**. EDPVR slope as an index of passive left ventricular chamber stiffness. **I**. Load-dependent cardiac work (LDCW), defined as the slope of the relationship between total PVA and end-systolic pressure. **J**. Preload-recruitable dP/dtmax, defined as the slope of the linear relationship between dP/dt_max_ and EDV as a load-independent index of myocardial contractility. Data are presented as box-and-whisker plots with individual data points overlaid; boxes represent the interquartile range (IQR, 25th–75th percentile), horizontal lines indicate the median, and whiskers extend to the minimum and maximum values (n=7-8 per group). All comparisons were performed by one-way ANOVA with Sidak’s multiple comparison test; exact p values are indicated between groups. Ctrl, naive healthy control mice; Sep, cecal slurry-induced septic mice administered vehicle; Sep+Ela, cecal slurry-induced septic mice administered elamipretide.

In addition to systolic impairment, indices of diastolic function were altered. The slope of the end diastolic pressure volume relationship (EDPVR) was increased in Sep mice, indicating impaired ventricular relaxation (**Fig. 2H**). Load-dependent cardiac work (LDCW), defined as the slope of the relationship between total PV area and end-systolic pressure, was also reduced, consistent with decreased EDV without changes in end-systolic volume (ESV), suggesting suboptimal preload conditions (**Fig. 2I, S2C-D**).

Ela treatment substantially improved these hemodynamic indices. In Sep mice, Ela increased SW, cardiac efficiency, Ees, and VAC, and restored load-independent indices of contractility, including PRSW and PRCW (**Fig. 2A-G**). Ela also improved diastolic function, as indicated by normalization of the EDPVR slope (**Fig. 2H**). Although LDCW tended to increase with Ela treatment, this effect did not reach statistical significance (P=0.056; **Fig. 2I**). Ela additionally improved cardiac reserve. The preload recruitable change in maximal pressure (PR dP/dt_max_), which was reduced in Sep mice, was restored with Ela treatment (**Fig. 2J**). Under hemodynamic stress (increased afterload), Ela further improved maximal cardiac functional capacity, mitigating sepsis-induced impairments in VAC and cardiac efficiency and restoring maximal generated pressure (Pmax) and theoretical maximal pressure (Piso) (**Fig. S2E-H**).

Collectively, these findings demonstrate that sepsis impaired intrinsic myocardial contractility, ventricular passive properties, and ventriculo-arterial coupling, and that Ela treatment improves these properties, resulting in more efficient cardiac energy utilization and improved cardiovascular performance.

### Sepsis induces cardiac lipidome remodeling that is partially attenuated by Ela

To evaluate changes in membrane lipid composition in the Sep heart and following Ela treatment, we performed untargeted lipidomic profiling of cardiac tissue at T12h, identifying 2,690 lipid features across Ctrl (n=6), Sep (n=7) and Sep+Ela (n=7) groups. PERMANOVA (Bray-Curtis dissimilarity) confirmed significant differences in the overall cardiac lipidome among groups (pseudo-F=1.95, P=0.022). Pairwise comparisons revealed significant separation between Ctrl and Sep (P=0.018) and between Sep and Sep+Ela (P=0.018), whereas Sep+Ela was not significantly different from Ctrl (P=0.24), suggesting partial normalization of the cardiac lipidome with Ela treatment (**Fig. 3A**). Comparison of Sep vs Ctrl groups revealed 40 differentially abundant lipid species at FDR<0.05, of which 17 were elevated and 23 were reduced in Sep hearts (purple circles, **Fig. 3B**). Comparison of Sep+Ela versus Sep animals identified 44 differentially abundant lipid species at FDR<0.05 (26 elevated, 18 reduced in Sep+Ela relative to Sep) (green circles; **Fig. 3B**). To characterize the biological properties of differentially abundant lipids, LION ontology enrichment analysis was performed using the full 2,690-feature detected lipidome as background (**Fig. 3C**). Although no LION terms survived FDR correction, several terms showed consistent nominal enrichment (P<0.05) across multiple analyses. The mitochondrial subcellular-localization term was enriched among sepsis-upregulated lipids, implicating mitochondria-associated lipid species as primary targets of sepsis-induced remodeling. Glycerophosphoglycerols (PG; GP04) were enriched specifically among sepsis-upregulated lipids, consistent with accumulation of cardiolipin precursor species in the septic heart.

**Figure 3.**
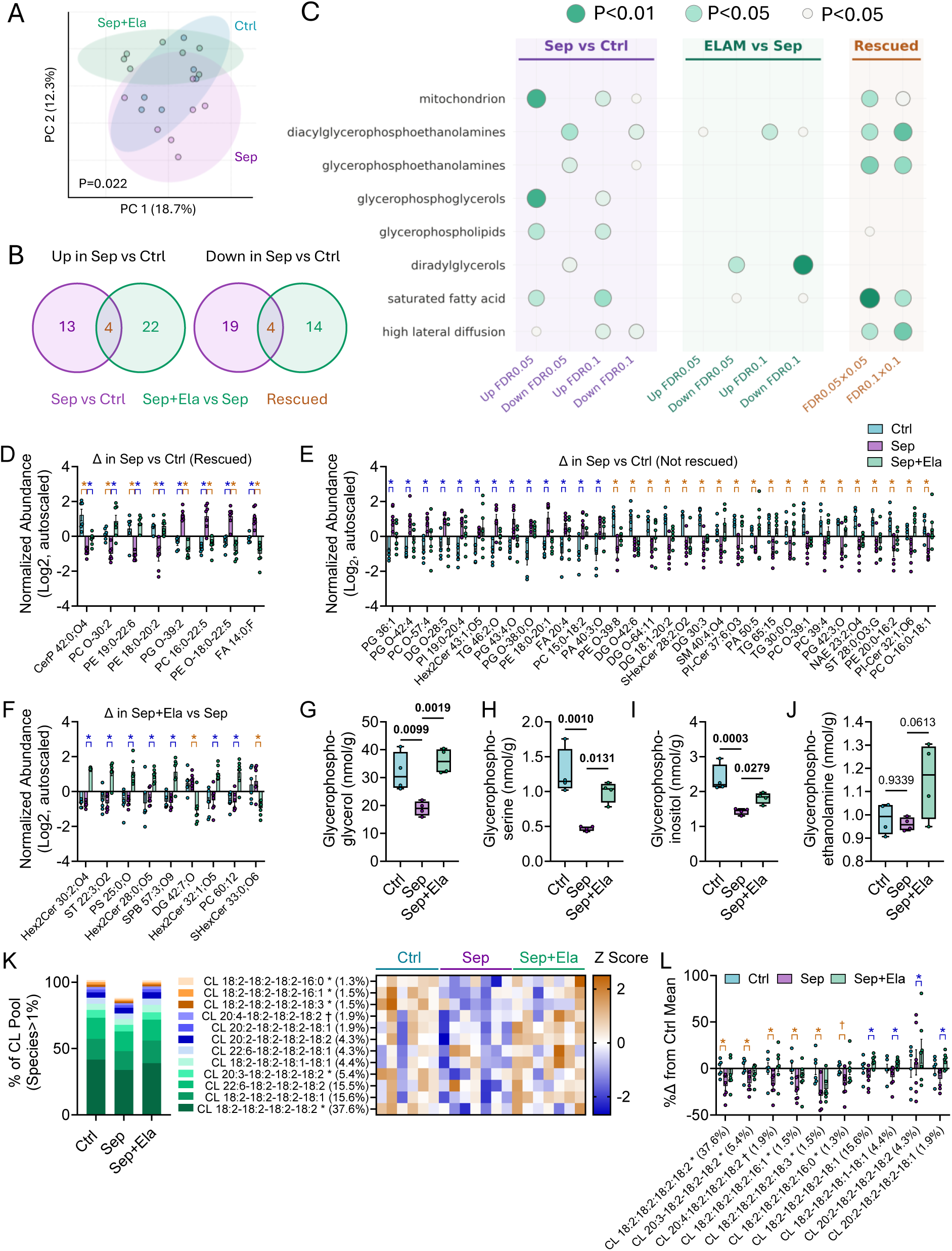
Elamipretide restores the cardiac lipidome in sepsis. **A.** Principal component analysis of the cardiac lipidome showing group separation among Ctrl, Sep, and Sep+Ela hearts (PERMANOVA, Bray-Curtis dissimilarity, P = 0.022). **B.** Bubble plot summarizing enrichment of lipid ontology terms across comparisons (Sep vs. Ctrl, Ela vs. Sep) at FDR thresholds of 0.05 and 0.1. Bubble size reflects the number of lipid species within each category; color intensity denotes enrichment significance. “Rescued” (rightmost column) denotes lipid categories that were perturbed in Sep vs. Ctrl and normalized in Sep+Ela. **C.** Venn diagrams depicting the overlap and unique differentially abundant lipid species between Sep vs. Ctrl and Sep+Ela vs. Sep comparisons, with “Rescued” species identified as those altered in sepsis and restored with Ela treatment. **D–F.** Individual lipid species significantly altered between groups, including **D**. rescued lipids, **E**. non-rescued lipids, denoting those perturbed in Sep vs. Ctrl but not normalized with Ela treatment, and **F**. lipids unchanged in Sep vs. Ctrl (P ≥ 0.5) but altered with Ela treatment. **G–J.** Box plots of select lipid derivative species across groups illustrating representative patterns of sepsis-induced perturbation and Ela-mediated restoration. **K.** Stacked bar plot of individual cardiolipin (CL) species composition across samples, showing the relative abundance of major CL molecular species and changes among treatment groups, and corresponding heatmap of CL species across Ctrl, Sep, and Sep+Ela groups. **L.** Comparison of individual CL species (that constitute >1% of total CL pool) between treatment groups. In panels **D–F** and **L**, blue asterisks denote significant increases, and orange asterisks significant decreases, relative to the reference group in each comparison (Ctrl for Sep vs. Ctrl; Sep for Sep+Ela vs. Sep; FDR < 0.05).

Cross-referencing the Sep vs Ctrl and Sep+Ela vs Sep comparisons identified 8 lipid species that were altered in Sep and restored toward control levels by Ela at FDR<0.05 in both analyses (**Fig. 3B-D**); the remaining 32 lipids altered in Sep were not affected by Ela (**Fig. 3E**). Of those rescued, 4 were elevated in Sep and reduced by Ela treatment (PG O-39:2, PC 16:0-22:5, PE O-18:0-22:5, FA 14:0;F), and 4 were reduced in Sep and restored by Ela (CerP 42:0;O4, PC O-30:2, PE 19:0-22:6, PE 18:0-20:2) (**Fig. 3D**). Notably, all 8 species showed directional reversal with Ela treatment; that is, none were altered in the same direction by both Sep and Ela, suggesting a specific lipid response to Ela. Among rescued lipids, LION analysis identified consistent nominal enrichment of mitochondrial localization, diacylglycerophosphoethanolamines, glycerophosphoethanolamines and saturated fatty acid terms, suggesting that Ela preferentially restores lipid species with mitochondria-associated functions and acyl chain composition with potential consequences for membrane biophysical properties including lateral diffusion (**Fig. 3C**).

Nine lipid species were significantly altered by Ela treatment despite showing no evidence of alteration in Sep versus Ctrl animals (i.e. FDR<0.05 for Sep+Ela vs Sep, and P≥0.5 for Sep vs Ctrl) (**Fig. 3F**). These elamipretide-specific lipids were dominated by glycosphingolipid species, including three hexosylceramide (Hex2Cer) species, a sulfatide (ST 22:3;O2), a sulfohexosylceramide (SHexCer 33:0;O6), and a sphingoid base (SPB 57:3;O9), with the remaining species comprising a phosphatidylserine (PS 25:0;O), a phosphatidylcholine (PC 60:12), and a diacylglycerol (DG 42:7;O). The predominance of sphingolipid species among Ela-specific changes is notable, as sphingolipids and cardiolipin share mitochondrial membrane microdomains and both contribute to mitochondrial membrane integrity and signaling. Whether these sphingolipid changes represent a direct secondary effect of cardiolipin stabilization, a compensatory membrane remodeling response, or an off-target effect of Ela treatment remains to be determined; nonetheless, these findings suggest that Ela’s lipidomic footprint extends beyond direct restoration of sepsis-altered species and may involve broader remodeling of the mitochondrial membrane sphingolipid landscape.

To validate these findings, we examined glycerophospholipid headgroup metabolites by targeted metabolomics. Glycerophosphoglycerol, glycerophosphoserine, and glycerophosphoinositol were all significantly reduced in septic hearts relative to controls and were partially restored by Ela-treatment (**Fig. 3G–I**), consistent with the class-level depletion of glycerophospholipid species identified by untargeted lipidomics. Notably, glycerophosphoethanolamine did not change with sepsis, but tended to increase in Sep+Ela mice relative to Sep (P=0.0613, **Fig. 3J**), mirroring the consistent enrichment of the glycerophosphoethanolamine ontology term among Ela-restored lipids identified by LION analysis and suggesting active remodeling of the PE pool by Ela independent of the sepsis-induced perturbations. Collectively, these targeted metabolomics findings corroborate the untargeted lipidomic data and reinforce the conclusion that sepsis broadly disrupts the cardiac glycerophospholipid landscape, and that elamipretide partially but consistently restores this disruption.

### Sepsis broadly depletes cardiac cardiolipin species, and Ela partially restores cardiolipin abundance

To directly assess the impact of Sep and Ela treatment on CL species, all 27 CL species detected in the untargeted lipidomic dataset were extracted and analyzed as a focused subset. Fractional pool composition was calculated based on individual species proportions of the summed raw intensity, revealing a non-uniform distribution dominated by three species, identified based on m/z match and concordance with published cardiac CL profiles^42^: CL 18:2-18:2-18:2-18:2 (37.6% of total CL intensity), CL 18:2-18:2-18:2-18:1 (15.6%), and CL 22:6-18:2-18:2-18:2 (15.5%) (**Fig. 3K**). Subsequent statistical analyses were restricted to the 12 species constituting >1% of the total CL pool, which collectively accounted for 95.2% of all detected CL. FDR correction was applied within this biologically defined subset. Of these, 11 (92%) were reduced in Sep hearts relative to controls, with 5 surviving FDR correction at q<0.05, and 6 at q<0.1 (**Fig. 3L**). The dominant species CL 18:2-18:2-18:2-18:2 was significantly reduced by 18.8% in Sep hearts (FDR=0.044). CL 20:3-18:2-18:2-18:2 (5.4% of pool; −15.8%; FDR=0.050), CL 18:2-18:2-18:2-16:1 (1.5%; −23.6%; FDR=0.031), CL 18:2-18:2-18:2-18:3 (1.5%; −29.3%; FDR=0.031), and CL 18:2-18:2-18:2-16:0 (1.3%; −17.9%; FDR=0.044) also survived FDR correction. The dominant species CL 18:2-18:2-18:2-18:2 accounted for 57% of the total pool-weighted CL deficit in Sep hearts, illustrating that the functional impact of CL depletion is disproportionately driven by changes in the most abundant species. By contrast, CL 16:0-20:4-18:2-18:3, which showed the greatest percent reduction at −29.3%, contributed only −0.43% weighted impact points owing to its 1.5% pool fraction.

Strikingly, all 12 species exceeding 1% of the total CL pool trended upward in Sep+Ela relative to Sep animals, indicating a uniform directional restoration of the CL pool by Ela (**Fig. 3L**). Four species survived FDR correction at P<0.05 for the Sep+Ela versus Sep comparison: CL 18:2-18:2-18:2-18:1 (15.6% of pool; +20.1%; FDR=0.013), CL 20:2-18:2-18:2-18:1 (1.9%; +22.1%; FDR=0.013), CL 18:2-18:2-18:1-18:1 (4.4%; +23.4%; FDR=0.024), and CL 20:2-18:2-18:2-18:2 (4.3%; +17.2%; FDR=0.024). One species, CL 20:4-18:2-18:2-18:2 (1.9% of pool) met formal rescue criteria with q<0.05 in both comparisons (−17.4% in Sep, p=0.045; +25.5% with Ela, p=0.026). The total pool-weighted Ela restoration was +14.3%, with the dominant species CL 18:2-18:2-18:2-18:2 contributing 40% of this incomplete restoration. Taken together, these findings provide evidence of broad CL depletion in the septic murine heart dominated by reductions in the most abundant CL species, with Ela inducing a uniform directional restoration of the CL pool.

### Elamipretide restores cardiac mitochondrial ultrastructure and function during sepsis

Given the established role of cardiolipin in maintaining cristae architecture and supporting electron transport chain function, we hypothesized that the sepsis-induced cardiolipin depletion would translate to structural and functional mitochondrial impairment. We therefore examined whether cardiac mitochondrial structure and respiratory capacity were compromised in Sep mice, and whether Ela treatment mitigated these deficits. TEM revealed marked ultrastructural changes in the hearts of Sep mice, characterized by regional cristae loss, disordered mitochondrial organization within the sarcomere, and disrupted outer and inner mitochondrial membranes, though this latter phenotype was variable (**Fig. 4A-B, S3A-E**). A marked accumulation of glycogen was also observed in Sep hearts, predominantly within intramyofibrillar rather than intermyofibrillar regions, as observed in Ctrl mice (**Fig. 4A, C**). Despite these abnormalities, mitophagic clearance was not regulated, based on the absence of autophagosomes in TEM images and lack of induction of *Pink1* and *Parkin* gene expression (**Fig. 4A**, S3B, F-G).

**Figure 4.**
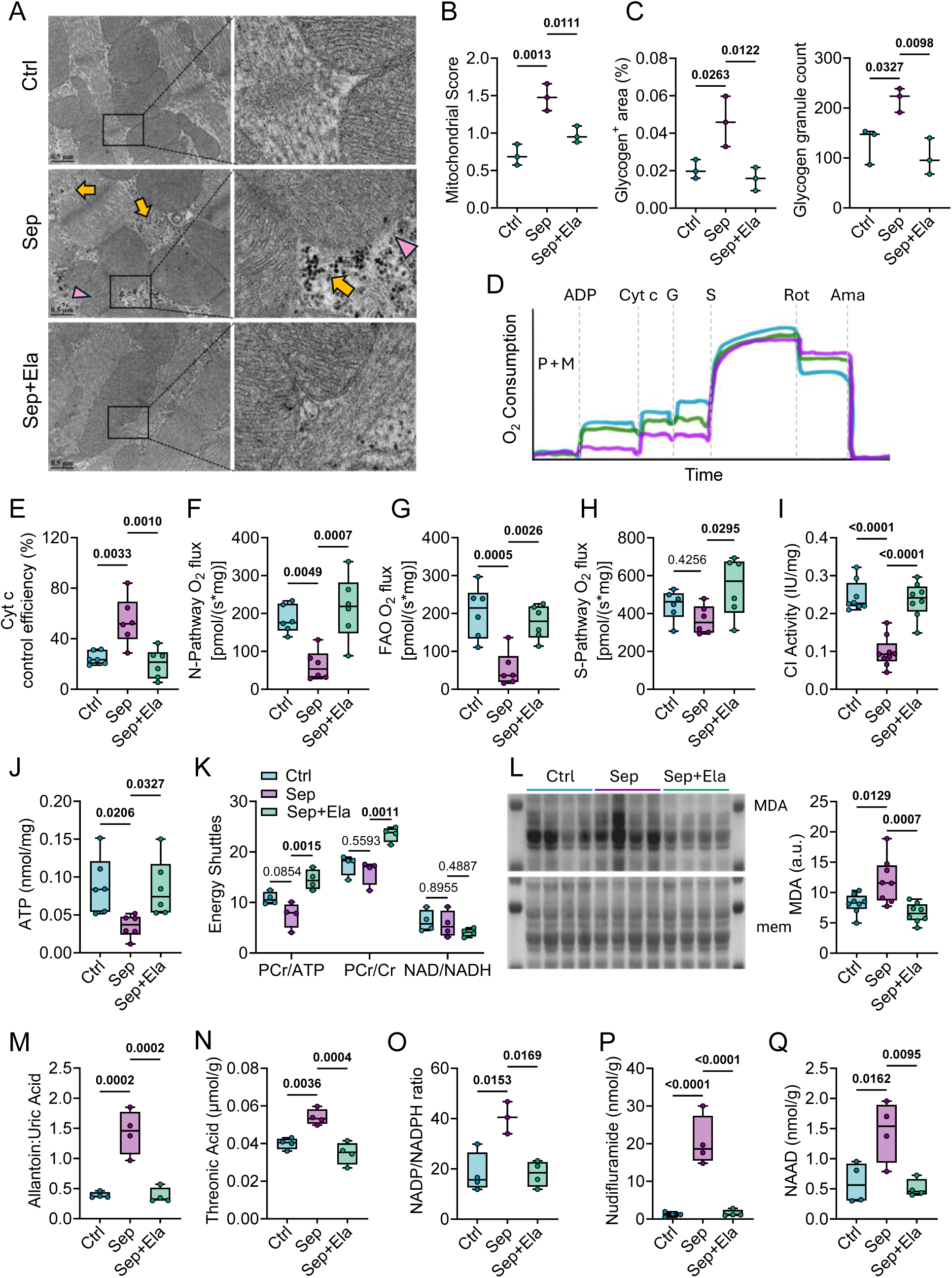
Sepsis causes mitochondrial ultrastructural abnormalities, impairs complex I–mediated respiration, and induces oxidative stress, effects that are alleviated by elamipretide. **A**. Representative transmission electron micrographs of left ventricle (LV) myocardium from Ctrl, Sep, and Sep+Ela mice. Yellow arrows indicate glycogen granule accumulation; pink arrowheads indicate disrupted mitochondrial membranes. Boxed regions are shown at higher magnification. **B**. Quantitative mitochondrial structural scoring. **C**. Glycogen-positive area (left) and glycogen granule counts (right) in LV myocardium (n=3 biological replicates; n=96-146 technical replicate images). **D**. Representative high-resolution respirometry traces of cardiac homogenates showing O₂ flux following sequential additions of pyruvate + malate (P+M), ADP, cytochrome *c* (Cyt *c*), glutamate (G), succinate (S), rotenone (Rot), and antimycin A (Ama). **E**. Cytochrome *c* control efficiency, reflecting mitochondrial membrane integrity. **F**. NADH (N)-linked pathway O₂ flux (through complex I) supported by pyruvate, malate, glutamate, ADP, and Cyt *c*. **G**. Fatty acid oxidation (FAO)–supported O₂ flux in the presence of palmitoylcarnitine, malate, ADP, and Cyt *c*. **H**. Succinate supported (S-pathway) O₂ flux (through complexes II) after inhibition with rotenone. **I**. Complex I enzymatic activity. **J**. Myocardial ATP content. **K**. Myocardial energy-state and redox ratios: phosphocreatine-to-ATP (PCr/ATP), phosphocreatine-to-creatine (PCr/Cr), NADP⁺/NADPH, and NAD⁺/NADH. **L**. Representative Western blot of malondialdehyde (MDA)-modified proteins (left) and corresponding densitometric quantification (right). **M**. Myocardial allantoin-to-uric acid ratio. **N**. Myocardial threonic acid concentration. **O.** Myocardial NAP^+^/NADPH ratio. **P**. Myocardial nudifloramide concentration. **Q**. Myocardial nicotinic acid adenine dinucleotide (NAAD) concentration. P-values were determined using one-way ANOVA followed by Sidak’s multiple-comparison test; n=3-6 per group. Ctrl = naive healthy control mice, Sep = cecal-slurry-induced septic mice receiving vehicle, Sep+Ela = cecal-slurry-induced septic mice receiving elamipretide. Data are presented as box-and-whisker plots showing the mean, interquartile range, minimum, and maximum values. Outliers were identified using Grubbs’ test (α = 0.05) and excluded prior to analysis.

Ultrastructural abnormalities were accompanied by marked impairments in mitochondrial functional capacity, assessed by high resolution respirometry of cardiac homogenates (**Fig. 4D-H**), despite the absence of changes in mitochondrial abundance (**S3H**). Leak O_2_ flux, measured in the presence of pyruvate and malate prior to ADP addition) did not differ among groups (data not shown). Cytochrome c control efficiency was significantly increased in Sep mice, indicating impaired outer mitochondrial membrane integrity (**Fig. 4E**). NADH-pathway (N-pathway; complexes I linked) O_2_ flux, supported by pyruvate, malate, glutamate, ADP and Cyt *c*, was significantly reduced in Sep mice compared with Ctrl mice (**Fig. 4F**). Similarly, fatty acid oxidation (FAO)-supported O_2_ flux, measured in the presence of palmitoylcarnitine, malate, ADP and Cyt *c*, was reduced in Sep hearts (**Fig. 4G**). In contrast, succinate-supported (S-pathway; complex II linked) O_2_ flux and complex IV alone were not altered by sepsis (**Fig. 4H, S3I**). ETS capacity, determined as the maxima O_2_ flux supported by convergent N- and S-linked pathway was significantly reduced in Sep mice (**Fig. S3J**). Collectively, these results indicate that septic hearts harbor intrinsic impairments in mitochondrial respiratory capacity. The concomitant reduction in N-pathway and FAO-supported O₂ flux, despite preserved S-pathway capacity, is consistent with a defect affecting electron supply to the Q-junction and/or complex I function. The reduction in complex I enzymatic activity (**Fig. 4I**) further supports a major contribution of complex I dysfunction to the respiratory phenotype observed in Sep hearts.

Functional respiratory deficits were associated with reduced myocardial ATP levels in Sep mice, though major energy shuttles were largely unaltered in Sep mice, as reflected by PCr/ATP, PCr/Cr and NAD/NADH ratios (**Fig. 4J-K**). Sep mice also exhibited evidence of cardiac oxidative stress, including increased myocardial malondialdehyde (MDA) levels, increased allantoin relative to uric acid, and elevated threonic acid, a product of ascorbic acid consumption (**Fig. 4L-P**). In addition, increased NADP:NADPH ratio and elevated NAAD levels in Sep hearts are consistent with increased ROS levels and metabolic stress (**Fig. 4K, Q**).

Ela treatment was associated with marked improvements in mitochondrial organization, fewer compromised membrane borders, and mitigated cristae loss, as well as reduced ectopic glycogen accumulation (**Fig. 4A-C, S3B-E**). The structural improvements were accompanied by a reduced Cyt c effect on respiratory capacity, prevention of impaired N-pathway and FAO mediated O_2_ flux, and restored ETS capacity (**Fig. 4D-G, S3J**), consistent with improved complex I activity (**Fig. 4I**). Interestingly, Ela treatment also increased S-pathway flux, despite no impairments caused by sepsis (**Fig. 4H**). Ela also restored LV free wall ATP levels and enhanced PCr/ATP and PCr/Cr ratios, suggesting improved energetic status and phosphocreatine reserve (**Fig. 4J-K**). Finally, Ela reduced MDA levels, normalized levels of oxidized derivatives, and reduced the NADP/NADPH ratio, consistent with reduced oxidative stress (**Fig. 4K-P**).

Together, these data indicate that sepsis causes mitochondrial structural anomalies and respiratory dysfunction, notably at complex I, resulting in bioenergetic impairment accompanied by evidence of oxidative stress, and that these pathologies are alleviated by Ela treatment.

### Murine Sepsis Perturbs Cardiac Substrate Utilization and Oxidative Metabolic Flux

Despite clear evidence of altered cardiac mitochondrial morphology, impaired respiratory capacity, and increased oxidative stress in Sep hearts, the observed deficits in cardiac contractile and relaxation performance could also reflect shifts in substrate utilization and metabolic flexibility. To address this possibility, we used an *ex vivo* working heart preparation to quantify glucose, fatty acids, and ketones.

Under baseline perfusion conditions (fatty acids, glucose and β-hydroxybutyrate present), Sep hearts exhibited lower cardiac work, developed pressure, and cardiac output, with a trend toward reduced cardiac efficiency, despite unchanged coronary flow, oxygen consumption or heart rate (**Fig. 5A-D, S4A-B**). These findings are consistent with the impaired contractility observed by echocardiography and PV loop experiments (**Fig. 1-2, S1-S2**). Ela treatment improved cardiac output and cardiac work, and produced a similar, although non-significant, trend toward increased developed pressure (**Fig. 5A-C)**. Ela also increased coronary flow relative to both Ctrl and Sep mice (**Fig. 5D**), whereas myocardial oxygen consumption (and heart rates) remained unchanged (**Fig. S4A-B**).

**Figure 5.**
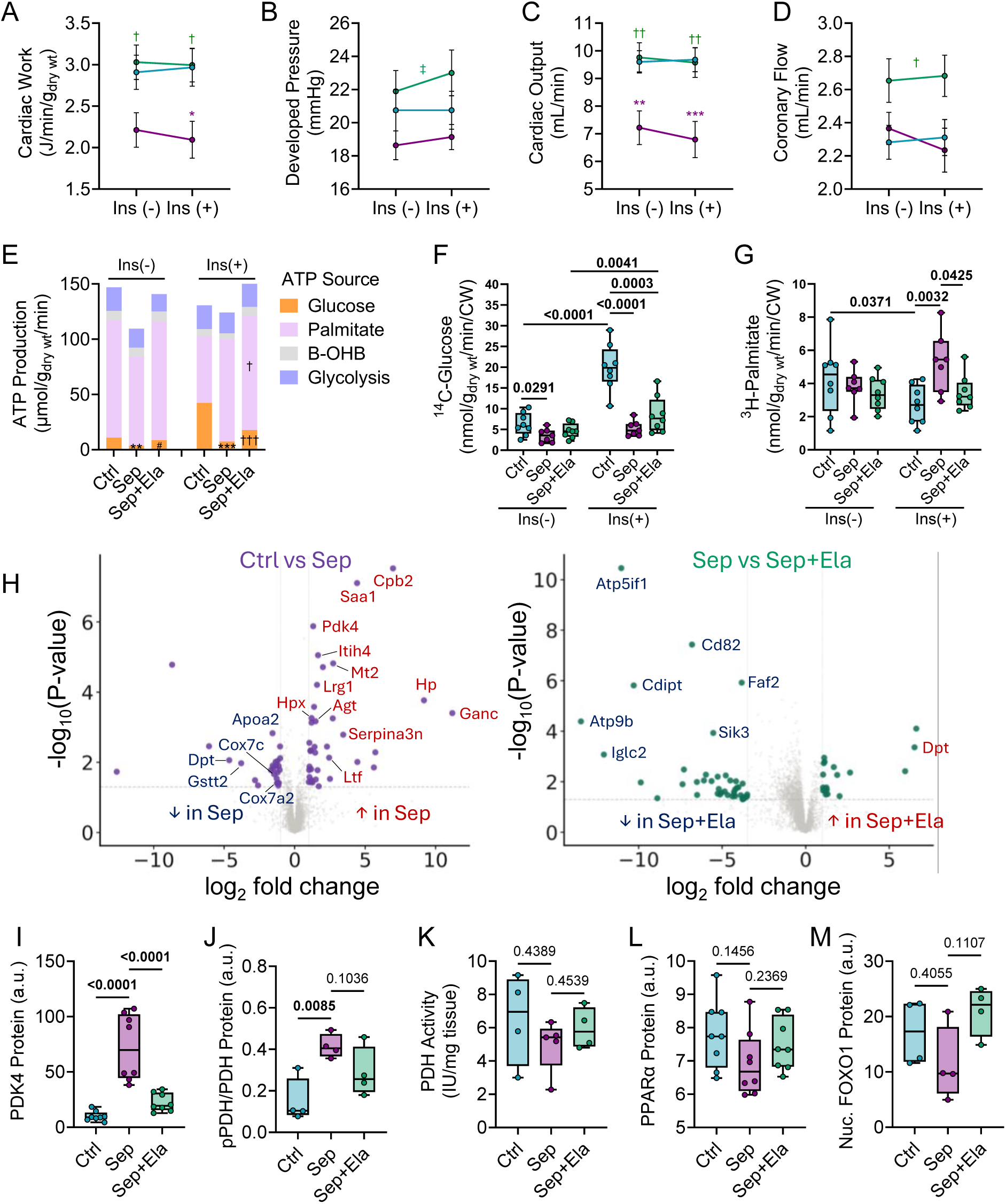
Sepsis impairs cardiac substrate oxidation and metabolic flexibility in the ex vivo working heart, and elamipretide enhances ATP production. A–D. Ex vivo cardiac functional performance measured in isolated working hearts across the span of 1 hour in the absence and presence of insulin (n=11-14). **A**. Cardiac work normalized to the dried heart mass, **B.** developed pressure, **C**. left ventricular cardiac output, and **D**. coronary flow. **E**. ATP production rates derived from glucose oxidation, glycolysis, palmitate oxidation, and β-hydroxybutyrate (βOHB) oxidation (n=6-8). *=Difference between Sep and Sep+Ela, #=Difference between Sep and Ctrl, †=Difference between Sep+Ela and Ctrl. **F**. Glucose oxidation rates determined from [U-^14^C]-glucose oxidation in the presence and absence of insulin and normalized to cardiac work (n=7-8). **G**. Palmitate oxidation rates determined from [9,10-^3^H]-palmitate oxidation in the presence and absence of insulin normalized to cardiac work (n=7-8). **H**. Volcano plots showing differentially expressed proteins in Ctrl vs Sep hearts (left), and Sep+Ela vs Sep hearts (right) (n=4). Representative Western blot measurement and corresponding densitometric quantification of **I**. Cardiac Pyruvate dehydrogenase kinase 4 (PDK4) protein abundance (n=8), **J**. Cardiac phosphorylated PDH-to-total pyruvate dehydrogenase (PDH) (n=4), **K**. Cardiac PDH activity (n=4), **L**. Cardiac PPARα protein abundance (n=8), **M**. Cardiac nuclear levels of FOXO1 protein. (n=4). Ctrl = naive healthy control mice, Sep = cecal-slurry induced septic mice receiving vehicle, Sep+Ela = cecal-slurry induced septic mice receiving elamipretide. Data are presented as box-and-whisker plots showing the mean, interquartile range, minimum, and maximum values. Statistical analyses were performed using one-way ANOVA followed by Sidak’s multiple-comparison test. Outliers were identified using the Grubbs’ test (α = 0.05) and excluded prior to analysis.

In the absence of insulin, palmitate was the dominant fuel source in all groups (**Fig. 5E**). Although glucose oxidation had a minor contribution to total ATP production under these conditions, glucose oxidation normalized to cardiac work was markedly reduced in Sep hearts compared to Ctrl hearts and was not restored Ela treatment (**Fig. 5F**). As expected, insulin administration increased glucose oxidation concomitant while suppressing fatty acid oxidation in Ctrl hearts, though no such metabolic shift occurred in Sep mice (**Fig. 5F-G**), suggesting cardiac insulin resistance.

Consistent with these metabolic alterations, unbiased proteomics analysis identified PDK4 as upregulated in Sep mice, a finding confirmed by Western Blot analysis (**Fig. 5H-I, S4A**). Because PDK4 phosphorylates and inhibits PDH, its induction provides a mechanistic explanation for the reductionin glucose oxidation. Accordingly, the phosphorylated-to-total PDH ratio was elevated in Sep hearts (**Fig. 5J, S4B**), whereas native PDH activity measured in cardiac homogenates under conditions that bypass regulatory phosphorylation remained unchanged (**Fig. 5K**), indicating that impaired pyruvate oxidation in sepsis resulted from regulatory inhibition of PDH rather than loss of intrinsic PDH catalytic capacity. Notably, increased PDK4 protein abundance occurred without corresponding changes in Pdk4 mRNA expression **(Fig. S4C),** suggesting post-transcriptional regulation. Cardiac PDK4 is a short-lived protein that is degraded by the mitochondrial Lon protease, with its stability governed by association with the PDH complex and modulated by fatty acid availability.^43^ The elevated fatty acid utilization observed in Sep hearts may therefore promote stabilization of PDK4 in its PDH-bound state, resulting in accumulation of active PDK4 protein and sustained inhibition of pyruvate oxidation. To further investigate potential upstream regulatory mechanisms, we examined signaling pathways previously implicated in PDK4 induction in genetic models of cardiolipin deficiency.^44^ Neither the canonical PPARα/PGC1α/PPARγ transcriptional axis nor nuclear FOXO1 abundance paralleled the changes in PDK4 induction (**Fig. 5L-M, S4A-C**), arguing against transcriptional activation through these regulators. Collectively, these findings support a model in which altered substrate utilization promotes post-translational PDK4 stabilization, contributing to its accumulation in septic hearts, whereas Ela reverses this process concomitantly with normalization of substrates metabolism.

Ela partially restored insulin responsiveness in Sep hearts, as evidenced by increased glucose oxidation **(Fig. 5F)**. However, this increase was modest and was not accompanied by a reduction in fatty acid oxidation (**Fig. 5G**). Similarly, the partial restoration of glucose oxidation mirrored the incomplete reduction in the p-PDH/PDH ratio with Ela despite robust normalization of PDK4 protein abundance (**Fig. 5J**), suggesting that additional regulatory mechanisms, including other PDK isoforms (PDK1, PDK2) and PDH phosphatase activity, contribute to the net phosphorylation state of PDH. Similar patterns were observed even when substrate oxidation rates were analyzed without normalization to cardiac work (**Fig. S4D-E**). Glycolytic flux, assessed by quantitative extraction of ^3^H from ^3^H-5-Glucose in the effluate, and ketone oxidative flux, measured by ^14^CO_2_ extraction, were not altered by Sep or with Ela treatment (**Fig. S4F-G**).

Taken together, while myocardial ATP output declined in Sep hearts concordant with blunted oxidative flux, Ela preserved oxidative metabolism, modestly enhanced glucose oxidation, and maintained fatty acid utilization, altogether augmenting ATP production (**Fig. 5E, S4H**).

### Sepsis is associated with profound cardiac metabolic dysregulation

We further examined whether the cardiac metabolic phenotype observed in working hearts was reflected in the *in vivo* cardiac metabolome. Notably, to establish whether the cardiac metabolic phenotype reflected altered systemic substrate availability or intrinsic cardiac dysfunction, we first quantified circulating substrate levels. Blood glucose, free fatty acids and ketone levels were reduced in Sep mice, and Ela did not alter these systemic levels (**Fig. 6A**), indicating that reduced substrate supply cannot account for the cardiac metabolic phenotype described below and that the effects of Ela on cardiac metabolism are not attributable to altered substrate availability.

**Figure 6.**
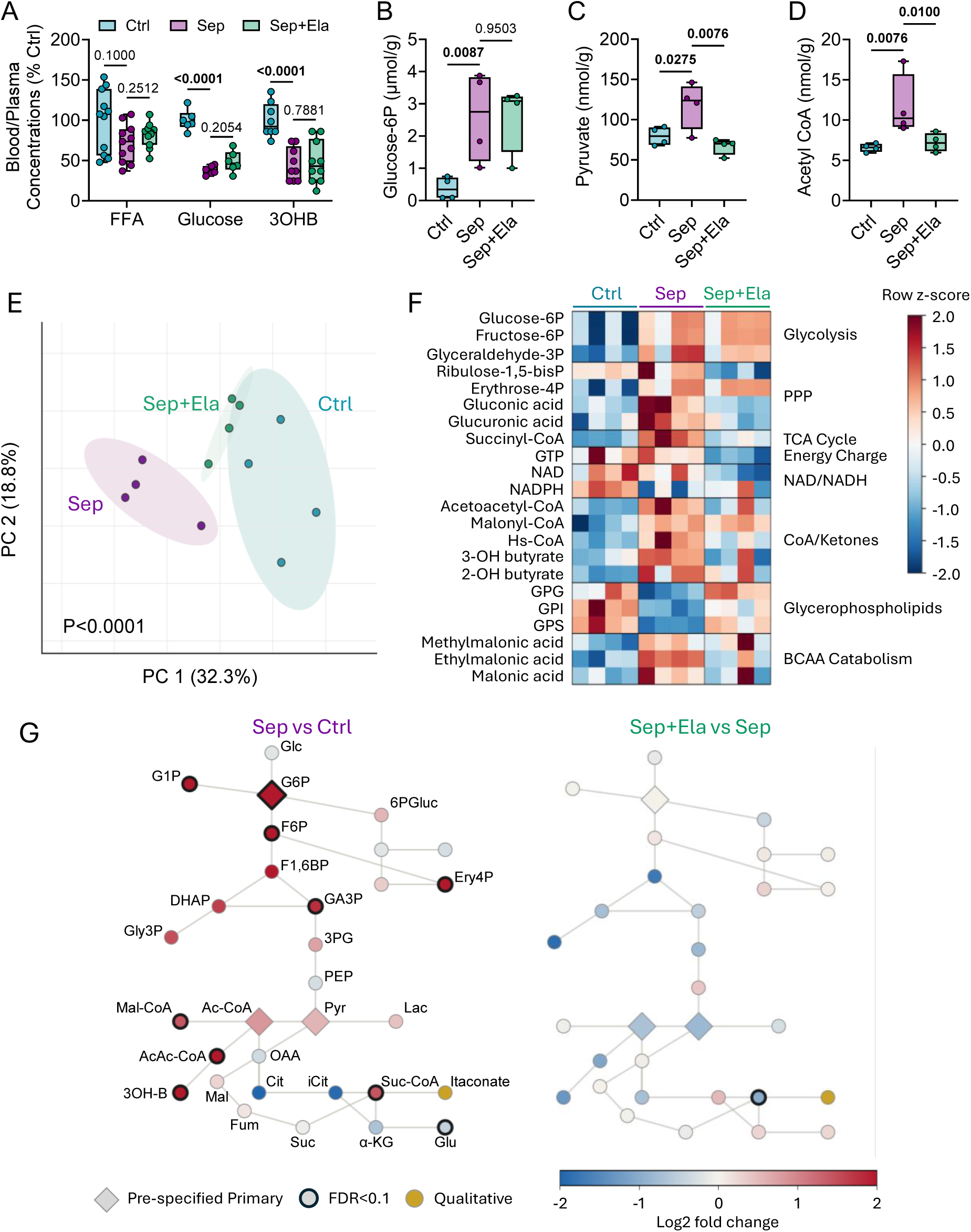
Cardiac metabolomic profiling reveals broad metabolic disruption in sepsis and elamipretide-mediated remodeling. **A**. Plasma levels of free fatty acids (FFA), glucose, and β-OHB. **B.** Myocardial glucose-6-phosphate levels. **C**. Myocardial pyruvate levels (n=4). **D**. Myocardial acetyl-CoA levels. **E**. Principal component analysis of metabolomic datasets across Ctrl, Sep, and Sep+Ela groups, displayed using PC2 (18.5% variance) and PC1 (32.3% variance) (n=4). **F**. Heat map generated from absolute concentrations of select myocardial metabolites. Each row was mean-centered to highlight within-metabolite variation, with a blue–white–red gradient denoting low, intermediate, and high relative concentrations, respectively (n=4). **G.** Metabolic network generated using the Shiny GATOM application and processed with the Escher program, based on absolute metabolite concentrations. Nodes represent individual metabolites, with edges indicating known biochemical connections. Node color reflects relative abundance from absolute concentrations, where red denotes higher levels, white indicates intermediate levels, and blue denotes lower levels (n=4). Ctrl = naive healthy control mice, Sep = cecal-slurry induced septic mice administered vehicle, Sep+Ela = cecal-slurry induced septic mice administered elamipretide. Data are presented as box-and-whisker plots showing the mean, interquartile range, minimum, and maximum values. Data were analyzed with one-way ANOVA with Sidak’s multiple comparison test. Outliers were identified and excluded using the Grubbs’ test with a significance level of α = 0.05.

The three pre-specified metabolomic primary endpoints supported the hypothesized pyruvate-oxidation bottleneck. Glucose 6-phosphate (6.8-fold; P=0.023), pyruvate (1.5-fold; P=0.028) and acetyl-CoA (1.8-fold; P=0.008) all accumulated in septic hearts **(Fig. 6B-D)**, concomitant with increased cardiac protein acetylation **(Fig.S5A**). Together, these findings are consistent with a mismatch between glycolytic substrate supply and downstream mitochondrial oxidative utilization. Elamipretide normalized pyruvate (P=0.008) and acetyl-CoA (P=0.010) toward control levels and reduced protein acetylation, whereas glucose 6-phosphate remained elevated (P=0.95). These data suggest that Ela relieved the downstream oxidative constraint associated with pyruvate oxidation while only partially resolving the accumulation of upstream glycolytic intermediates (**Fig. 6B-D, S5A**).

Across the broader exploratory panel, the cardiac central carbon metabolome separated by group, with PCA distinguishing Ctrl, Sep, and Sep+Ela hearts (PERMANOVA P<0.0001; **Fig. 6E**). Sep hearts diverged from Ctrl along the principal axis, while Ela-treated hearts were displaced toward the control distribution, indicating partial normalization of the global metabolic profile. A total of 26 metabolites differed between Sep and Ctrl hearts (FDR<0.1), with changes that clustered coherently by pathway (**Fig. 6F**). Glycolytic intermediates upstream of pyruvate accumulated as a block (fructose 6-phosphate, glyceraldehyde 3-phosphate, glucose 1-phosphate; all FDR < 0.1), reinforcing the concept of glycolytic congestion seen in the primary endpoints. The non-enzymatic glucose oxidation products gluconate and glucuronate were also elevated (each FDR < 0.1), consistent with the combined effects of glycolytic substrate accumulation and the increased oxidative milieu in septic hearts.

The most pronounced exploratory changes involved the CoA-ester and ketone pools. Succinyl-CoA showed the strongest individual change of any metabolite (2.7-fold increase; FDR=0.027), accompanied by accumulation of acetoacetyl-CoA (4.6-fold; FDR=0.070), malonyl-CoA (2.7-fold; FDR=0.095), free CoA (4-fold; FDR=0.095), and ketone bodies 3-hydroxybutyrate (5.5-fold; FDR=0.075) and 2-hydroxybutyrate (3.9-fold; FDR=0.095) (**Fig. 6F-G, S5B-G**). This coordinated expansion is consistent with impaired downstream mitochondrial oxidative utilization, resulting in accumulation of acetyl-CoA and related CoA esters, together with increased myocardial ketone body abundance. SCOT abundance was unchanged in septic heart (**Fig. S5H**), indicating that altered ketone handling cannot be attributed to reduced expression of this key ketolytic enzyme. The expanded acetyl-CoA pool likewise does not appear to reflect enhanced β-oxidation. CPT1β, the rate limiting enzyme of mitochondrial fatty acid entry, was downregulated at the protein level (**Fig. S5I**); while malonyl CoA, an endogenous CPT1 inhibitor, was elevated (**Fig. S5D**). In addition, ethylmalonate (ethylmalonic acid, 3.6-fold, FDR = 0.070), a non-canonical product of butyryl-CoA metabolism indicative of impaired fatty acid breakdown, and methylmalonic acid (1.5-fold, FDR = 0.075) a byproduct of odd-chain fatty acid metabolism, accumulated in septic hearts (**Fig. S5I-K**). Together, these findings are consistent with constrained mitochondrial substrate utilization, resulting in the accumulation of acetyl CoA and ketone bodies despite reduced systemic substrate availability.

In contrast, citrate and isocitrate exhibited downward trends in sepsic hearts (FDR = 0.34 and 0.24, respectively), while glutamate was significantly reduced (FDR=0.095). This pattern is consistent with reduced entry of carbon into the TCA cycle and diminished oxidative metabolism. Membrane-associated glycerophospholipid species (glycerophosphoinositol, -serine, and -glycerol) were coordinately decreased (all FDR≤0.075), paralleling the cardiolipin remodeling identified in the lipidomic analysis (**Fig. 3)**. Itaconate, an inflammation-induced immunometabolite, was undetectable in all control hearts but present in all septic hearts, revealing a metabolic signature of inflammation that was absent under basal conditions (**Fig. S5L**).

### Elamipretide normalizes the septic metabolic signature

Elamipretide reversed a substantial portion of the septic metabolic phenotype: 12 metabolites differed between Sep+Ela and Sep hearts at FDR < 0.1, with the CoA-ester, ketone, and pyruvate changes shifting back toward control values (**Fig. 6F**), consistent with improved cardiac mitochondrial oxidative substrate utilization. When mapped within the metabolic pathway network, the broad accumulation of central carbon metabolites observed in Sep hearts **Fig. 6G, left**) was attenuated with Ela (**Fig. 6G, right**), indicating coordinated normalization of cardiac metabolic homeostasis rather than isolated changes in individual metabolites. The persistence of glycolytic intermediate accumulation despite Ela treatment (**Fig. 6F-G**) indicates that normalization of the septic metabolic phenotype was incomplete. Nevertheless, the selective reduction in pyruvate, acetyl-CoA, ketone bodies, and related CoA-esters suggests improved downstream mitochondrial substrate utilization, consistent with Ela’s proposed action on the inner mitochondrial membrane.

### Ela prevents mortality and residual cardiac dysfunction in Sep mice

To investigate post-sepsis recovery, mice were treated with antibiotics and fluids from T12h to T84h, and survival was assessed until T30d (**Fig. 7A**). Sep mice exhibited increased mortality over the subsequent 30 days, with most deaths occurring between 10 and 20 days post-inoculation. Ela treatment (at T1h) completely prevented these late deaths (**Fig. 7B**).

**Figure 7.**
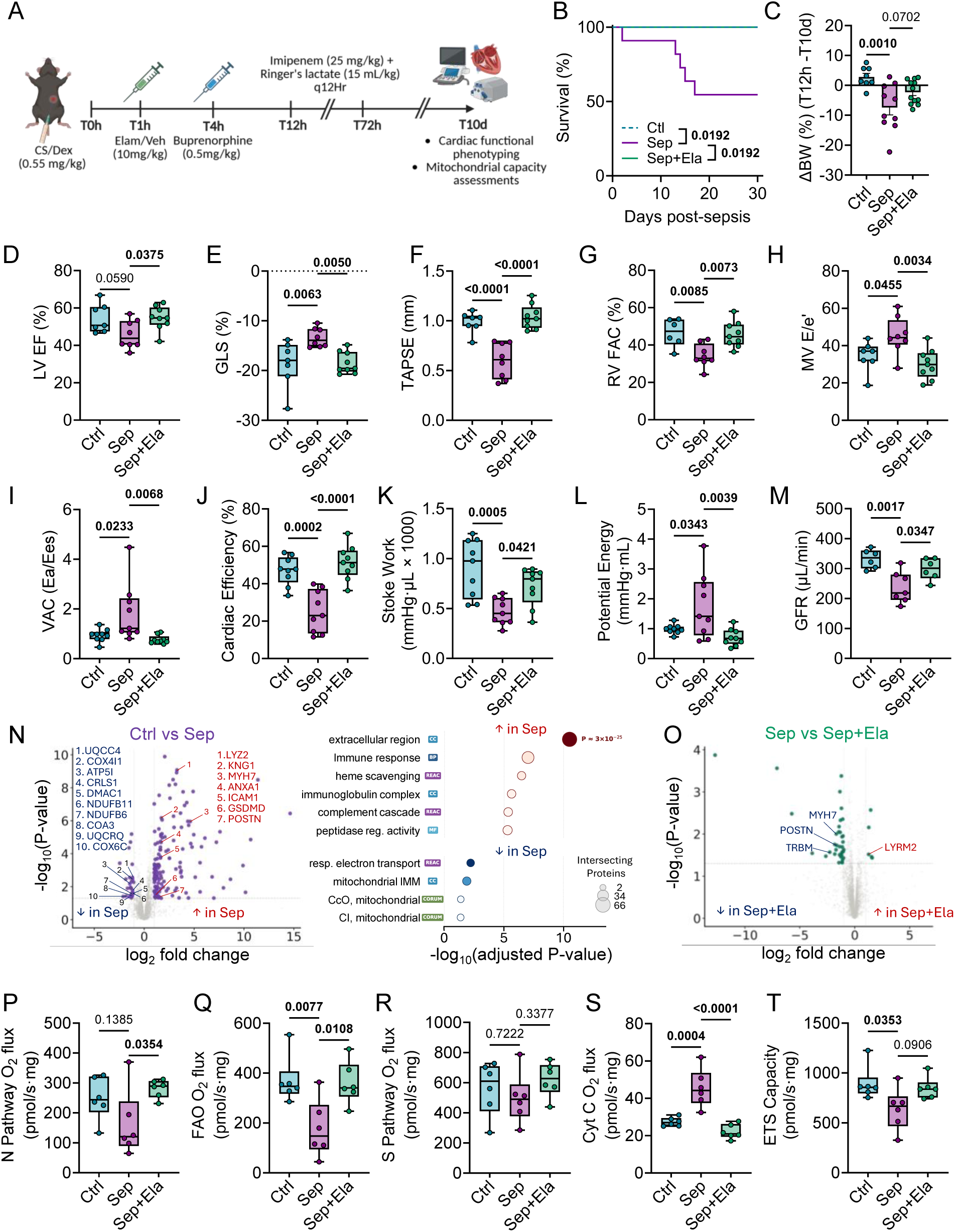
Ela prevents Persistent cardiac and mitochondrial impairments after sepsis recovery. **A**. Experimental timeline outlining sepsis induction and the ensuing sepsis recovery protocol, and subsequent post-sepsis assessments. **B**. 30-day survival rates post-sepsis recovery. Survival differences between groups were assessed using the Log-rank (Mantel-Cox) test (n=11-14). **C**. Bodyweight change from 12hr post-sepsis induction to 10 days post-sepsis recovery (n=8-12). **D**. Left ventricular ejection fraction (LV EF) assessed with echocardiography (n=7-9). **E**. Global longitudinal strain (GLS) measured with echocardiography (n=7-9). **F**. Tricuspid annular plane systolic excursion (TAPSE) measured with echocardiography (n=7-9). **G**. Right ventricular fractional area change (FAC). **H**. Mitral valve E/e’ ratio as an indicator of diastolic function measured with echocardiography. **I**. Ventriculo-arterial coupling (VAC). **J**. Cardiac efficiency and **K**. stroke work determined with pressure-volume loops (n=9). **L**. Potential energy measured with pressure-volume loops (n=7-9). **M**. *In vivo* glomerular filtration rate (GFR) determined transdermally normalized to bodyweight (n=6-7). **N-O.** Volcano plots showcasing differentially expressed proteins between **N**. Ctrl vs Sep 10 days post-sepsis (n=4) with pathway enrichment analysis of salient processes altered post-sepsis (right) and **O**. differentially expressed proteins between Sep+Ela vs Sep 10 days post-sepsis (n=4). **P**. Increases in oxygen flux in response to cytochrome C administration, representing replenishment of cytochrome C loss assessed with mitochondrial respirometry (n=6). **Q-T**. Oxygen flux measured by high resolution respirometry in the presence of **Q**. pyruvate, glutamate and malate, showing respiration through complex I, or **R**. palmitoylcarnitine and malate, showing fatty acid oxidation, or **S**. Complex II mediated oxygen consumption in the presence of pyruvate and malate with addition of rotenone (n=6). **T**. Respirometry-derived maximal electron transport capacity in response to uncoupler 2,4-dinitrophenol (n=6). Ctrl = naive healthy control mice, Sep = cecal-slurry induced septic mice administered vehicle, Sep+Ela = cecal-slurry induced septic mice administered elamipretide. Data are presented as box-and-whisker plots showing the mean, interquartile range, minimum, and maximum values. Data were analyzed with one-way ANOVA with Sidak’s multiple comparison test. Outliers were identified and excluded using the Grubbs’ test with a significance level of α = 0.05.

Because most sepsis-related mortality occurred between days 10 and 20, we selected day 10 as the cardiovascular assessment timepoint to characterize cardiac and molecular phenotype before the period of maximal sepsis-related mortality, thus minimizing survivor bias (**Fig. 7A**). At this time point, circulating cytokines remained elevated in Sep mice (**Fig. S7A-D**) but were attenuated compared to T12h (**Fig. 1C, S1B**), indicating at least partial resolution of systemic inflammation. As in the acute phase, Ela treatment had no effect on circulating cytokines in Sep mice (**Fig. S7A-D**). However, Sep mice exhibited the greatest weight loss compared to other groups (**Fig. 7C**), consistent with greater systemic burden of illness. Echocardiography at T10d revealed persistent cardiac dysfunction in Sep compared to naïve Ctrl mice. Although EF was not significantly different between groups (**Fig. 7D**), Sep mice had reduced GLS, CO, SV, (**Fig. 7E**, **S7E-F**), as well as reduced TAPSE, RV FAC (**Fig. 7F-G**), and MV E/e’ (**Fig. 7H**), indicating residual impairments in LV and RV systolic function and LV diastolic function. In contrast, some parameters, such as GCS, were no longer different in Sep mice (**Fig. S7G**), suggesting partial recovery of cardiac function. PV loop assessments corroborated these findings: VAC was impaired, primarily due to alteration in Ees rather than Ea (**Fig.7I, S7H**), and residual intrinsic contractility deficits were evident from reduced PRSW, PRCW, and PRdP/dt_max_ (**Fig. S7I-K**), though no differences in LDCW were observed (**Fig S7L**). Cardiac efficiency remained decreased in Sep hearts, associated with a reduction in stroke work with a concomitant increase in potential energy, consistent with reduced mechanical efficiency characterized by decreased stroke work and increased potential energy (**Fig. 7K-M**).

Ela treatment at T1h improved cardiac function at T10d. Echocardiography revealed improvements in LV systolic (**Fig. 7D-E, S7G**), RV systolic function (**Fig. 7F-G**), and LV diastolic function (**Fig. 7H**). PV loop analysis confirmed that Ela was associated with improved contractility (Ees), VAC and cardiac efficiency compared to untreated Sep mice (**Fig. 7I-L, S7H**). Ela treatment was also associated with improved renal function (GFR, **Fig. 7M**). Notably, there were no overt changes in cardiac mass (HW/TL), LV chamber volumes (EDV), or LV wall thickness (LV AWd and LV PWd) between Sep and Sep+Ela groups (**Fig. S7M-P**), suggesting subcellular deficits, rather than large-scale morphological remodeling, underlie the functional impairments observed.

We next investigated the metabolic underpinnings of post-sepsis cardiac dysfunction. Proteomic analyses of cardiac tissue collected at T10d revealed persistent downregulation of mitochondrial proteins in Sep compared to Ctrl mice, including ETS components: complex I subunits (NDUFB6, NDUFB11, and NDUFB3), complex IV subunits (COX6C, COX2, COX4I1), complex III and V proteins (UQCRQ, ATP5I, ATP5F1C), and ETS assembly factors (UQCC4, PET100) (**Fig. 7N**). Cardiolipin synthase (CRLS1) was also downregulated, suggesting impaired mitochondrial membrane remodeling and reduced capacity to support ETS integrity. Conversely, Sep hearts exhibited enrichment of proteins involved in inflammation and cell death (ANXA1, complement components C3 and C4B, GSDMD, BCL2L1, STAT1, and HMGB2), stress response (SOD3, CPB2), and structural remodeling (SORBS1, TMEM33, and creatine kinase M-type CKM) compared to Ctrl. In Sep+Ela hearts, fewer mitochondrial proteins were downregulated relative to Sep hearts, though some deficits persisted, including subunits of complex I (NDUFA1, NDUFB11), complex IV (COX2, COX7R, COX16), ATP synthase subunit (ATP5F1C), mitochondrial ribosomal protein RM16, and the mitochondrial assembly factor DMAC1 (**Fig. 7O**). These findings suggest that Ela mitigated the extent of mitochondrial protein loss following sepsis, particularly within the ETS.

Finally, we sought to determine whether residual cardiac dysfunction coincided with persistent impairments in mitochondrial function. Consistent with the reductions in ETS components, O_2_ flux remained reduced in the presence of N-pathway substrates (pyruvate, malate, glutamate; Fig. 7P), and following addition of palmitoylcarnitine (**Fig. 7Q**), but not in the presence of succinate and rotenone (**Fig. 7R**), altogether indicating persistent defects affecting NADH-linked respiratory capacity and localized, at least in part, to complex I. These defects in mitochondrial capacity were prevented with Ela treatment (**Fig 7P-R**). Notably, CS activity and CIV activity were not different among groups (**Fig. S7Q-R**), suggesting complex-specific impairments rather than reduced whole-mitochondrial content. Moreover, increased O_2_ flux with cytochrome c in Sep mice indicates persistent defects in mitochondrial membrane integrity, and this was fully restored with Ela treatment (**Fig. 7S**). Maximal ETS capacity (flux uncoupled from ATP synthase) was also reduced in Sep mice, with a trend toward restoration with Ela that did not reach statistical significance (**Fig. 7T**).

## Discussion

SICD is associated with increased short-term mortality,^45,46^ yet it remains underrecognized as a distinct clinical entity. Without a clear understanding of the underlying pathophysiology, targeted treatments for SICD cannot be realized. Here, using a well-characterized murine model of SICD, we performed a detailed functional and metabolic characterization, including direct assessments of metabolic flux in the heart, and identify metabolic modulation as an intervention strategy for both acute and residual cardiac dysfunction in sepsis.

### Elamipretide improves cardiac function independent of immunomodulation

SICD is characterized by global (biventricular) dysfunction encompassing systolic and diastolic impairments,^47^ all of which were evident in Sep mice. Hypokinetic LV wall motion and poor ventricular strain in this murine model resemble the wall-motion abnormalities observed in septic patients.^48^ Ela-mediated improvements in LV and RV systolic performance translated into systemic benefits, including improved cardiorenal hemodynamics, partial recovery of glomerular filtration rate, and reduced pulmonary edema. Diastolic dysfunction, attributable in part to ATP depletion impairing active calcium-reuptake and relaxation kinetics, and the progressive systolic depression that accompanies bioenergetic failure, were both alleviated by elamipretide. Ventriculo-arterial uncoupling in sepsis reflects intrinsic weakening of ventricular contractile function even in the presence of reduced afterload, and the associated decline in cardiac efficiency, which was restored by Ela, suggested altered myocardial energetics and substrate utilization. Importantly, comparable upregulation of inflammatory cytokines and immunometabolites such as itaconate in Sep and Sep+Ela groups indicates that Ela’s functional benefits are independent of systemic or myocardial immunomodulation. These observations prompted us to examine whether these changes were paralleled, or driven, by myocardial mitochondrial and metabolic remodeling.

### Cardiolipin loss underlies mitochondrial structural and functional deterioration

CL deficits are observed in human hypertrophic and dilated cardiomyopathies and are implicated in metabolic and functional deterioration.^21^ In sepsis, dysregulation of membrane lipid composition involving CL provided the therapeutic rationale for Ela, which stabilizes mitochondrial membranes and supports electron transport.^29^ CL is essential to the function of ETS complexes and is a target for peroxidative damage, owing to its localization on the inner mitochondrial membrane (IMM), proximal to sites of ROS generation, and its content of polyunsaturated fatty acyl chains. Our findings reveal widespread reductions in multiple CL species during sepsis, accompanied by mitochondrial respiratory dysfunction that likely underlies the observed metabolic and functional decline.

Ela-mediated improvements in CL levels were accompanied by partial restoration of mitochondrial morphology, increased respiration, elevated ATP levels, and reduced ROS and lipid peroxidation, consistent with outcomes reported in a canine model of advanced heart failure.^49^ In sepsis, Ela may preserve CL levels through direct biophysical interactions that protect CL from peroxidative damage, support cristae network integrity, and thereby mitigate ROS generation by improving ETS function. Consistent with this, reduced MDA-adduct formation implies that Ela limits ROS-mediated peroxidation of CL and other phospholipid acyl chains, protecting mitochondrial membrane integrity.

Notably, studies of Ela in ischemia-reperfusion injury,^29^ chronic heart failure,^30^ and Barth syndrome^31^ have generally reported reductions in CL content that were not restored by Ela treatment. In contrast, our results reveal that Ela not only restored the abundance of multiple CL species but also increased levels of several phosphatidyl subclasses (PC, PE, PS, PI) enriched in acyl chains that may contribute to CL or phospholipid remodeling. The IMM is particularly enriched with PEs (and their precursor PS), PC, and CL, all of which were broadly perturbed in septic hearts and restored with Ela. Perturbations in multiple PG species were also evident in sepsis, with restoration following Ela treatment; given that PG is a key intermediate in CL biosynthesis, preservation of PG pools may support CL synthesis or remodeling capacity.^50^ Supporting inter-phospholipid exchange of acyl chains, evidence indicates that the availability of phospholipid acyl chains is a primary determinant of CL composition.^51^ Phospholipid composition, specifically the length and saturation of acyl groups, dictates cristae curvature and bioenergetic capacity^52^ and loss of phospholipid diversity may contribute to membrane permeability in sepsis. Moreover, levels of glycerophosphoglycerol (GPG)and glycerophosphoserine GPS, headgroup metabolites integral to CL structure and overall membrane lipid architecture, were disrupted in Sep and improved in Sep+Ela hearts, suggesting preserved membrane lipid homeostasis.

The lipidomic analysis presented here provides evidence that sepsis-induced cardiomyopathy is accompanied by profound remodeling of the cardiac mitochondrial membrane lipidome, and that treatment with the cardiolipin-stabilizing peptide elamipretide partially restores this disruption. The consistent enrichment of PE and PG species, the two major phospholipid classes of the inner mitochondrial membrane, across multiple analytical comparisons points to a coordinated disruption of mitochondrial membrane lipid homeostasis in the septic heart. PE interacts directly with cardiolipin to support the structural integrity and functional efficiency of the electron transport chain, and its depletion in septic cardiac tissue is consistent with the mitochondrial dysfunction that characterizes sepsis-induced cardiomyopathy. The observed accumulation of PG species, as the immediate biosynthetic precursor to cardiolipin, may reflect impaired conversion of PG to CL by cardiolipin synthase – a bottleneck previously described under conditions of oxidative stress.^53^

Direct quantification of 27 cardiac CL species provided further support for this model and revealed that the impact of sepsis on the CL pool is disproportionately driven by changes in the most abundant species. Analysis was focused on the 12 species accounting for 95.2% of the total CL pool, 11 of which trended downward in sepsis and 5 of which survived FDR correction. The 18.8% reduction of the dominant species (CL 18:2-18:2-18:2-18:2) alone contributed 57% of the total pool-weighted CL deficit. Critically, all 12 biologically significant CL species trended upward in elamipretide-treated animals, representing a uniform directional restoration of the cardiac CL pool that would not have been apparent from percentage change data alone. This finding is consistent with Ela’s known mechanism of selectively binding and stabilizing cardiolipin at the inner mitochondrial membrane and provides direct lipidomic evidence that the drug engages its intended target in the septic heart.

The partial reversal of both PE depletion and PG accumulation by elamipretide, a peptide that binds selectively to CL and stabilizes IMM architecture, provides mechanistic support for cardiolipin as a central mediator of lipid remodeling in septic cardiomyopathy. The biophysical change among differentially abundant lipids further suggests that sepsis shifts the cardiac inner mitochondrial membrane toward a more fluid state, a change that would be expected to impair the function of the respiratory chain complexes and reduce OXPHOS efficiency.

### Elamipretide restores complex I function and attenuates oxidative stress

The respiratory phenotype observed in septic hearts suggests that cardiolipin disruption preferentially compromises mitochondrial pathways that depend on efficient electron transfer through complex I. Although fatty acid oxidation was also impaired, the preservation of succinate-supported respiration argues against a generalized collapse of the electron transfer system and instead points toward a defect affecting complex I-dependent oxidative metabolism. This interpretation is strengthened by the reduction in complex I enzymatic activity and by the coordinated loss of several complex I subunits identified by proteomic analysis. Restoration of respiratory function by Ela is therefore consistent with preservation of cardiolipin-dependent complex I function rather than a nonspecific enhancement of mitochondrial respiration.^54^ Complex I is a primary source of cellular superoxide production; impaired electron flux at this site is conducive to oxidative stress. Ela may attenuate oxidative stress by restoring complex I function and alleviating stalled electron flow. Concordantly, Chatfield et al. demonstrated that elamipretide selectively improved complex I activity (without affecting Complex II) in isolated mitochondria from explanted failing human hearts, without altering CL levels, suggesting that Ela stabilizes cardiolipin-protein interactions critical for individual complex function.

Reduced complex I activity has been linked to CL content in ischemia-reperfusion injury,^29^ a condition characterized by oxidative stress similar to sepsis; notably, supplementation with exogenous CL restored complex I function while peroxidized CL did not, indicating that peroxidation impairs the functional role of CL.^55^ Whereas earlier work proposed that Ela’s benefits may involve assembly of respiratory supercomplexes, their functional significance remains contested, with recent work demonstrating that impaired supercomplex assembly does not invariably compromise mitochondrial respiratory capacity.^56^ Our data are most consistent with a model in which Ela preserves complex I-dependent mitochondrial function through CL stabilization. We posit that Ela preserves membrane integrity via CL interactions, thereby curtailing electron leak and ROS generation from the ETS.

### Sepsis uncouples glycolysis from oxidative metabolism, and elamipretide partially restores metabolic flexibility

While the septic heart exhibits metabolic inflexibility,^57,58^ augmenting mitochondrial respiratory capacity can enhance oxidative flux and substrate metabolism to support cardiac function. Reduced basal and insulin-stimulated glucose utilization in Sep hearts reflects an intrinsic impairment in glucose oxidation, likely compounded by the pronounced hypoglycemia of sepsis, which limits availability in vivo. Elevated PDK4, increased oxidative stress, and impaired OXPHOS collectively compromise the septic heart’s capacity to oxidatively metabolize glucose and reinforce an insulin-resistant phenotype. This disruption creates a metabolic bottleneck: impaired glucose oxidative flux promotes glycogen accumulation that pervades the contractile apparatus and may hinder sarcomere mechanical function.^59^ Accordingly, intrinsic contractile capacity was undermined in sepsis, as denoted by decreased PRSW and Ees.

Despite having no effect on blood glucose, Ela improved glucose oxidation, suggesting at least partial preservation of metabolic flexibility. Though the measured effects appear modest, glucose oxidative flux derived from exogenously supplied ^14^C-glucose may underestimate total glucose oxidation in Ela-treated hearts, as it does not account for oxidation of unlabeled endogenous substrates, consistent with reduced accumulation of glycogen stores with Ela. The preservation of glycolytic flux despite reduced glucose oxidation in sepsis suggests uncoupling of glycolysis from mitochondrial oxidative metabolism. Ela improved downstream oxidative substrate utilization, as evidenced by normalization of pyruvate and acetyl-CoA levels, although accumulation of upstream glycolytic intermediates persisted.

### PDK4 induction in sepsis is post-translationally regulated and decoupled from canonical transcriptional drivers

The elevated PDK4 protein and PDH phospho-inhibition observed in septic hearts identify PDK4 as a central regulatory node restricting cardiac glucose oxidation. Although AMPK–FOXO1–PDK4 signaling has been described as the upstream driver of PDK4 induction in genetic models of cardiolipin deficiency (Barth syndrome),^44^ this axis was not engaged in our model: nuclear FOXO1 was not elevated in septic hearts and the canonical PGC1α/PPAR transcriptional program moved inversely with PDK4. Critically, Pdk4 mRNA was not significantly elevated in sepsis despite the marked increase in PDK4 protein, indicating that the upregulation in this setting is predominantly post-translational. Cardiac PDK4 has an exceptionally short half-life (∼1h) and is specifically degraded by the mitochondrial Lon protease, with stability governed by its association with the PDH complex and modulated by fatty acid availability.^43^ We propose that the elevated cardiac fatty acid utilization observed in septic hearts may stabilizes PDK4 in its PDH-bound, active state, allowing protein accumulation without transcriptional induction. Ela, by restoring mitochondrial substrate flexibility and normalizing fatty acid over-reliance, removes the substrate-dependent stabilization signal and permits Lon-mediated turnover, explaining why PDK4 protein falls with treatment despite persistent systemic inflammation. This mechanism reframes PDK4 in sepsis as a downstream readout of mitochondrial metabolic state rather than as a primary transcriptional driver of metabolic inflexibility and distinguishes sepsis-induced cardiac dysfunction from the chronic genetic CL-deficiency context in which the AMPK–FOXO1 axis dominates. It also provides a mechanistic explanation for the partial nature of glucose-oxidation recovery: reduction of PDK4 protein relieves one regulatory constraint on PDH, but other contributors (PDK1, PDK2, and PDH phosphatase activity) and persistent inflammatory drive likely account for the incomplete normalization of p-PDH and pyruvate oxidation in elamipretide-treated septic hearts.

### Impaired ETS function drives a downstream metabolic bottleneck affecting multiple substrate pathways

In parallel with impaired glucose oxidation, fatty acid oxidation remained the principal contributor to residual oxidative flux in the beating septic heart. Even in the presence of insulin, fatty acid oxidation remained elevated when normalized to cardiac work. However, sole reliance on fatty acid oxidation is energetically inefficient compared to glucose oxidation^60–62^ and during sepsis, fatty acid-derived ATP production may be insufficient to meet cardiac energy demands, potentially compromising performance. Despite this reliance on fatty acid oxidation, elevated ethylmalonate, linked to impaired acyl-CoA dehydrogenase activity, indicated disrupted β-oxidation and incomplete substrate processing, consistent with partial dysfunction of this pathway. Ela-mediated attenuation of ethylmalonate accumulation, together with upregulation of CPT1β and increased acetylcarnitine (Car 2:0) levels, suggests preservation of β-oxidation processes even in the wake of sepsis-induced suppression of fatty acid oxidative enzymes.

Collectively, these findings suggest that sepsis disrupts the integration of substrate oxidation with mitochondrial energy transduction, leading to accumulation of metabolic intermediates across multiple pathways. Acetyl-CoA, succinyl-CoA, and ketone body precursors all accumulated in septic hearts despite reduced circulating β-hydroxybutyrate, consistent with impaired mitochondrial substrate utilization rather than altered substrate supply. Accumulation of methylmalonate and ethylmalonate, byproducts of impaired odd-chain and short-chain fatty acid oxidation respectively, further supports the presence of constrained mitochondrial throughput. Ela restored complex I function and re-established forward electron flux, with downstream normalization of these accumulated intermediates. Consistent with this, Ela reduced cardiac acetyl-CoA and succinyl-CoA levels, attenuated ethylmalonate accumulation, and normalized byproducts of odd-chain fatty acid metabolism, without necessarily increasing β-oxidation rates per se, but rather improving substrate throughput.

### Sepsis disrupts cardiac redox and polyamine homeostasis

Beyond substrate metabolism, sepsis induced perturbations in redox-sensitive and stress associated metabolic pathways. Elevated levels of nudifloramide (2PY, an end-product of NAD^+^ catabolism) and NAAD indicate accelerated NAD^+^ turnover and altered NAD^+^ biosynthesis. Despite a stable NAD^+^/NADH ratio, an elevated NADP^+^/NADPH ratio indicates depletion of the antioxidant NADPH pool and engagement of antioxidant defenses, consistent with the oxidative stress evidence in septic hearts. Ela normalized myocardial levels of nudifloramide and NAAD, indicating restoration of NAD turnover.

### Cardiac metabolic signatures in sepsis mirror clinical biomarkers of poor prognosis

Intriguingly, a clinico-metabolomic analysis correlating plasma metabolites with survival outcomes in sepsis patients delineated metabolic profiles of non-survivors that mirror several perturbations observed in our septic murine hearts. Elevated serum allantoin, acetyl-CoA, pyruvate, urea, and glucuronate were particularly associated with non-survival, as was hyperacetylation of amino acids (N-acetylalanine, N-acetylthreonine).^63^ Additionally, acetylcarnitine levels were reduced and gluconate levels increased in sepsis compared to uninfected patients.^63^ These circulating metabolic profiles are consistent with impaired mitochondrial substrate utilization across tissues, and the convergence of these signatures in the heart, as observed here, supports the possibility that cardiac metabolic injury contributes to poor survival. While NT-proBNP and NGAL remain standard markers of cardiac and tissue injury, there is a compelling case for incorporating metabolite-based markers that may provide a more nuanced indication of cardiac function and prognosis in SICD.

### Post-sepsis cardiomyopathy persists despite resolution of inflammation

In the CIP model, decrements in GLS revealed underlying cardiac dysfunction post-sepsis. GLS is a sensitive indicator of ventricular function and has superior prognostic utility compared to ejection fraction across heart failure subtypes; abnormal LV GLS during sepsis predicted increased major adverse cardiac events 2 years post-sepsis.^48^. Elevated myocardial Myh7 (β-MHC) levels at day 10 further supports the presence of post-sepsis cardiomyopathy despite the absence of cardiac hypertrophy, reflecting adverse ultrastructural remodeling. Impaired preload recruitable stroke work (PRSW), a measure of intrinsic contractility, was accompanied by significant metabolic compromise: marked downregulation of ETS subunits and persistent reductions in complex I protein abundance, complex I-dependent respiratory capacity, and complex I enzymatic activity. Elevated periostin and decorin, indicative of myofibroblast activation and extracellular matrix remodeling, suggest emerging maladaptive pro-fibrotic remodeling. Although our characterization spanned the acute (T12h) and recovery (T10d) phases of SICD in this murine model, many of the observed metabolic and molecular alterations resemble those seen in chronic heart failure and inherited cardiomyopathies, highlighting the severity and clinical relevance of SICD.

## Conclusions

Although sepsis imposes a range of cellular stressors, these ultimately converge on shared pathways that disrupt cardiac metabolism, which is a hallmark feature of SICD. Reduced mitochondrial respiratory reserve and impaired complex I-dependent oxidative metabolism emerge as key mechanisms contributing to both acute and persistent cardiac dysfunction. Specifically, we propose that impaired electron transfer creates a central metabolic bottleneck associated with accumulation of glycolytic intermediates, CoA esters, ketone bodies, and markers of incomplete substrate oxidation, ultimately promoting bioenergetic failure, oxidative stress, and maladaptive metabolic remodeling. By stabilizing cardiolipin and preserving mitochondrial membrane integrity, elamipretide restores oxidative metabolism, improve substrate utilization, and attenuates oxidative stress, thereby mitigating both acute and residual cardiac dysfunction. This positions metabolic modulation as a promising therapeutic avenue to address both acute and residual cardiovascular burden of SICD. Given that cardiolipin disruption and complex I dysfunction are common features of multiple forms of cardiac and mitochondrial disease, these findings support metabolic modulation as a promising therapeutic strategy not only in SICD but potentially across a broader spectrum of disorders characterized by impaired mitochondrial energy transduction

## Author contributions

Conceptualization, J.V., S.L.B., K.M., G.L.; Methodology, J.V., S.L.B., G.L., J.U., H.L., A.D., K.M.; Investigation, J.V. C.W., C.H., A.W., T.B., M.L., I.K., S.N.L.; Formal Analysis, J.V., SL.B.; Data Curation and Visualization, J.V., SL.B.; Writing – Original Draft, J.V., S.L.B.; Writing – Review & Editing, all authors; Funding Acquisition, S.L.B., J.V.; Supervision, S.L.B., G.L., J.U.

## Data Availability

The datasets generated during this study are available from the first and corresponding author upon reasonable request.

