## Supplementary material for "Mitochondrial-targeted therapy with elamipretide preserves cardiac function and prevents late mortality in murine polymicrobial sepsis": Data supplement

**Data supplement for:**

**Supplementary Table S1: Primer Sequences**

| Gene ID | Forward Primer (5'→3') | Reverse Primer (5'→3') |
|---|---|---|
| *Pink1* | GCACAACATCCTTGTGGAGTGG | CATTGCCACCACGCTCTACACT |
| *Parkin* | CCAGAGGAAAGTCACCTGCGAA | GTTCGAGCAGTGAGTCGCAATC |
| *Pgc1α* | AGCCGTGACCACTGACAACGAG | GCTGCATGGTTCTGAGTGCTAAG |
| *Pdk4* | TCGAACTCTTCAAGAATGCC | GGTCAGTAATCCTCAGAGGAACA |
| *Pparα* | GCAATGGCTTTATCACACG | CCGATCTCCACAGCAAATTA |
| *Pparγ* | GTACTGTCGGTTTCAGAAGTGCC | ATCTCCGCCAACAGCTTCTCCT |

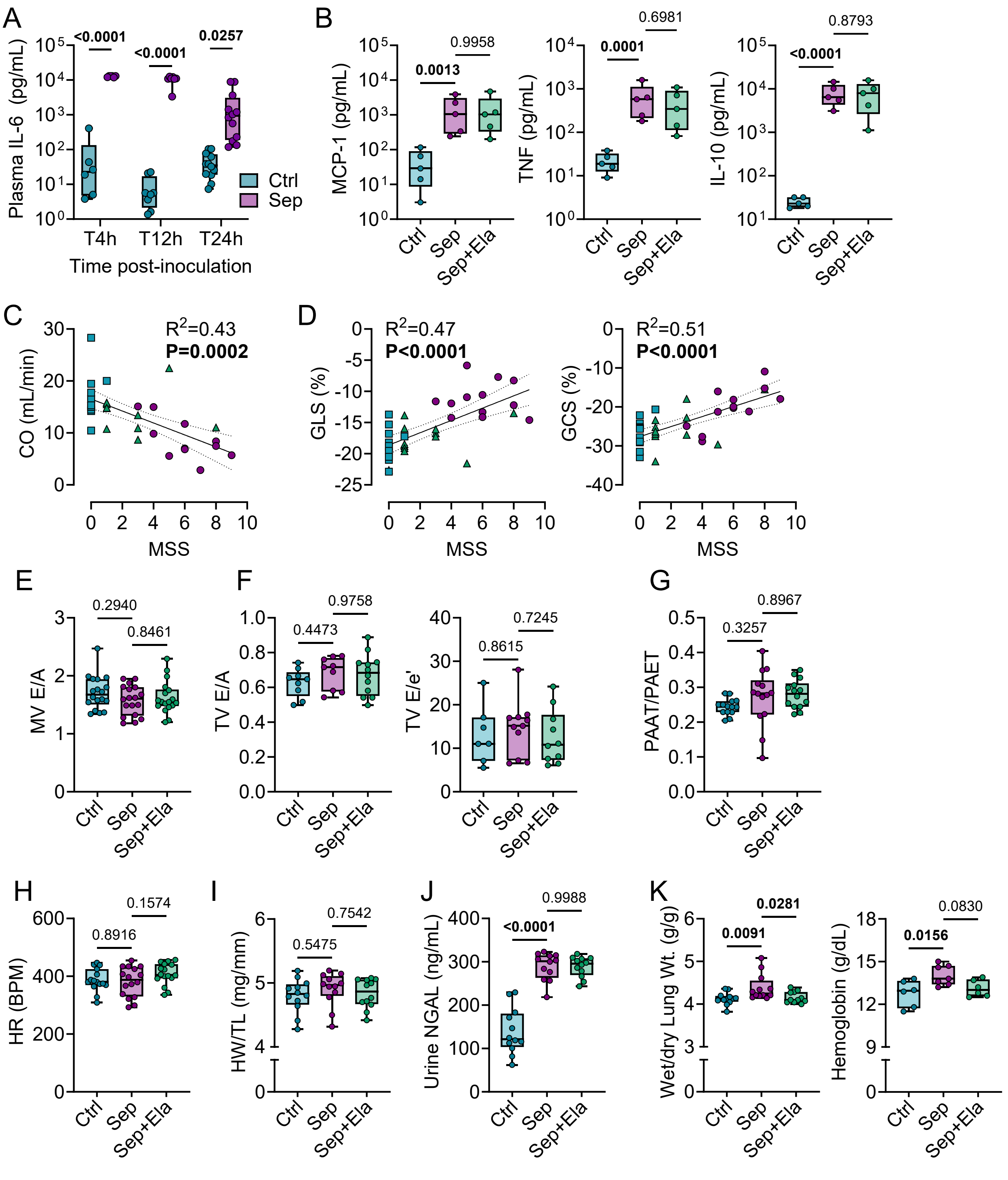


**Figure S1. Inflammatory cytokine profiles, additional echocardiographic parameters, and markers of end-organ injury. A**. Plasma interleukin-6 (IL-6) concentrations in Control and Septic mice at T4h, T12h, and T24h following cecal slurry inoculation (n=3–6 per timepoint). **B**. Plasma concentrations of monocyte chemoattractant protein-1 (MCP-1), tumor necrosis factor-α (TNF-α), and interleukin-10 (IL-10) at T12h (log-transformed; n=5). **C**. Correlation of cardiac output (CO) with murine sepsis score (MSS) at T12h. **D**. Correlations of global longitudinal strain (GLS; left) and global circumferential strain (GCS; right) with MSS at T12h. **E**. Mitral valve E/A ratio assessed by Doppler echocardiography (n=18). **F**. Tricuspid valve E/A ratio and E/e' ratio (n=7–12). **G**. Ratio of pulmonary artery acceleration time to pulmonary artery ejection time (PAAT/PAET) as an index of pulmonary vascular resistance (n=14). **H**. Heart rate (BPM, beats per minute). **I**. Heart weight normalized to tibia length (HW/TL; n=12). **J**. Urine neutrophil gelatinase-associated lipocalin (NGAL) concentrations (n=12). **K**. Wet-to-dry lung weight ratio (left; n=11–12) and hemoglobin concentration (right; n=6–7). Data are presented as box-and-whisker plots with individual data points overlaid; boxes represent the interquartile range (IQR, 25th–75th percentile), horizontal lines indicate the median, maximum and minimum values. All comparisons were performed by one-way ANOVA with Sidak's multiple comparison test; exact p values are indicated between groups. Ctrl, naive healthy control mice; Sep, cecal slurry-induced septic mice administered vehicle; Sep+Ela, cecal slurry-induced septic mice administered elamipretide.

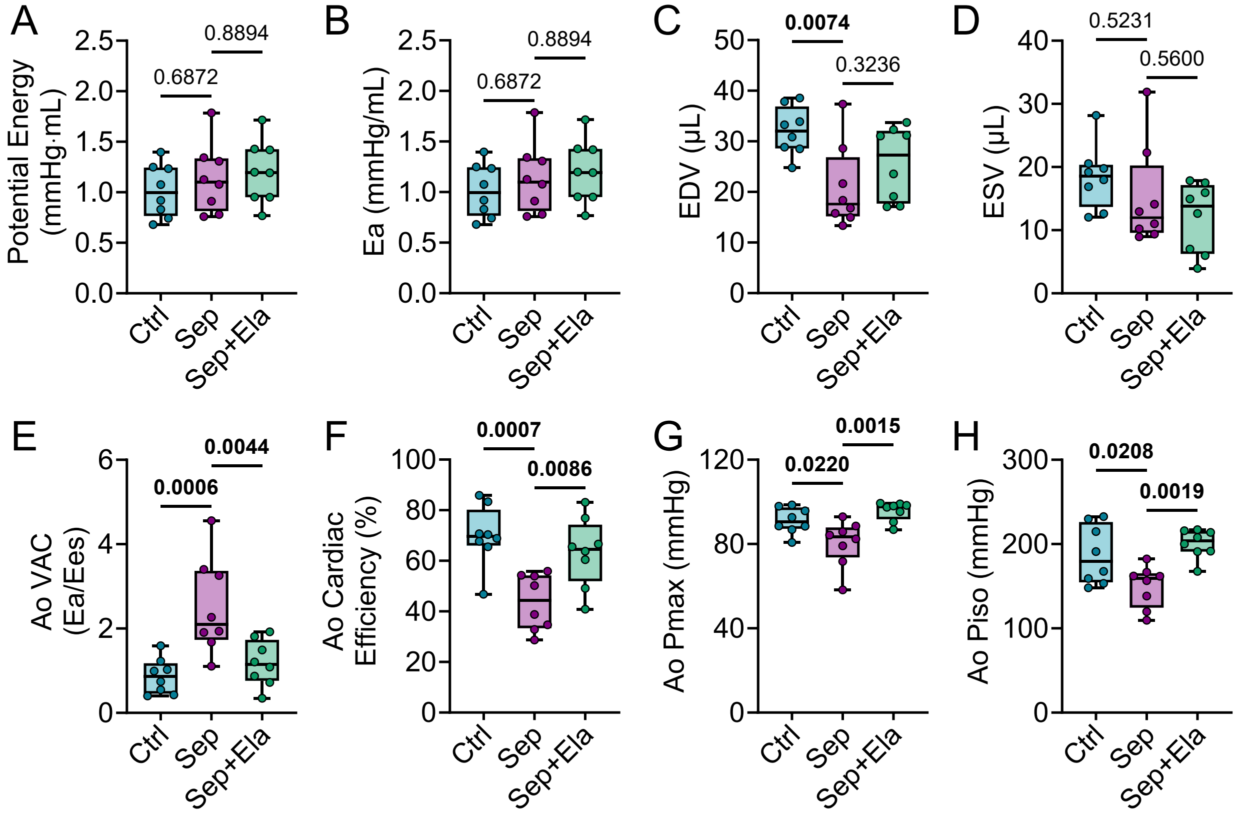


**Figure S2. Pressure-volume loop-derived indices of contractile capacity and reserve assessed at baseline and during aortic occlusion.** **A**. Potential energy, representing the region bounded by the ESPVR, EDPVR, and the isovolumetric relaxation line. **B**. Arterial elastance (Ea). **C**. End-diastolic volume (EDV). **D**. End-systolic volume (ESV). **E**. Ventriculo-arterial coupling (VAC) and **F**. cardiac efficiency assessed during aortic occlusion (AO) as indices of contractile reserve. **G**. Left ventricular maximal pressure (Pmax) generated and **H**. estimated theoretical maximal isovolumetric pressure (Piso) by the left ventricle, both assessed during aortic occlusion. Data are presented as box-and-whisker plots with individual data points overlaid; boxes represent the interquartile range (IQR, 25th–75th percentile), horizontal lines indicate the median, and whiskers extend to the minimum and maximum values (n=8 per group for all panels). All comparisons were performed by one-way ANOVA with Sidak's multiple comparison test; exact p values are indicated between groups. Ctrl, naive healthy control mice; Sep, cecal slurry-induced septic mice administered vehicle; Sep+Ela, cecal slurry-induced septic mice administered elamipretide.

A

| Mitochondrial Criteria | 0 | 1 | 2 |
|---|---|---|---|
| Cristae loss | Less than 10% of mito have a patch of cristae loss | Between 10-50% of mito have patches of cristae loss | More than half of the mito have patches of cristae loss |
| Membrane (OMM/IMM) integrity | Less than 10% of mito have blurry membranes | Between 10-50% of mito have blurry membranes | More than half of mito have blurry membranes |
| Sarcomere alignment | Mostly straight sarcomeric alignment | Regional variations – some sections misaligned | Very disordered alignment |

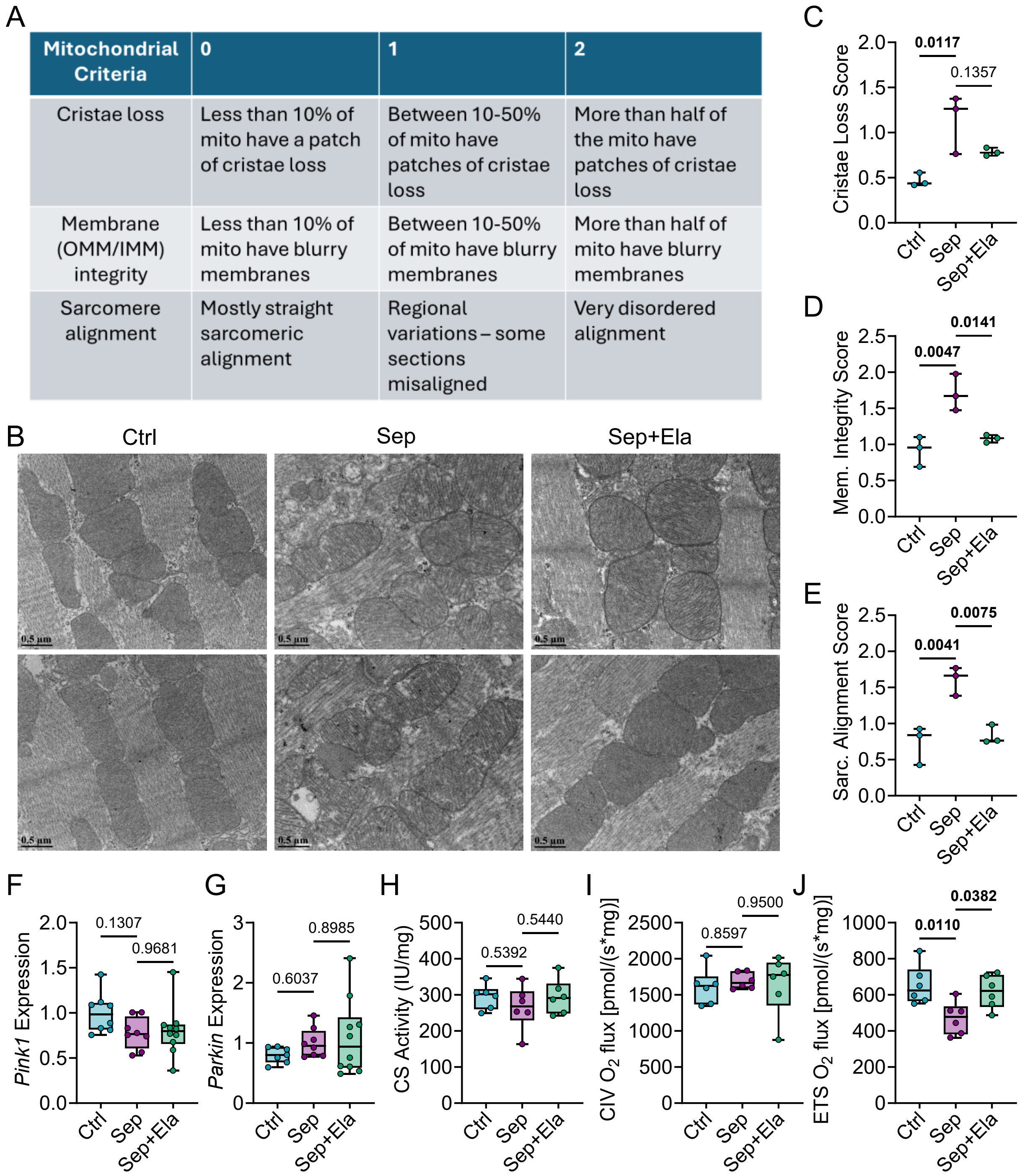


**Figure S3. Mitochondrial ultrastructural scoring criteria and detailed assessments of cardiac mitochondrial morphology, glycogen distribution, mitophagy, mitochondrial content, and respiratory capacity. A**. Scoring rubric for semi-quantitative assessment of mitochondrial ultrastructure by TEM. Three criteria, including cristae loss, outer and inner mitochondrial membrane (OMM/IMM) integrity, and sarcomere alignment, were each scored from 0 (minimal disruption) to 2 (severe disruption). **B**. Representative TEM micrographs of hearts. **C**. Individual mitochondrial cristae loss scoring. **D**. Mitochondrial membrane integrity scores. **E**. Sarcomere alignment scores, reflecting the organization of mitochondria in relation to myofibril orientation. **F-G**. Myocardial *Pink1* and *Parkin* gene expression. **H.** Citrate synthase activity, as a marker of mitochondrial content. **I**. Complex IV activity. **J**. Electron transport system (ETS) capacity, determined as maximal $O_2$ flux supported by convergent NADH- and succinate-linked pathways. P-values shown determined using one-way ANOVA followed by Sidak's post hoc test; n = 3-6 per group. Ctrl = naive healthy control mice, Sep = cecal-slurry-induced septic mice receiving vehicle, Sep+Ela = cecal-slurry-induced septic mice receiving elamipretide. Data are presented as box-and-whisker plots showing the mean, interquartile range, minimum, and maximum values. Outliers were identified using Grubbs' test ($\alpha = 0.05$) and excluded prior to analysis.

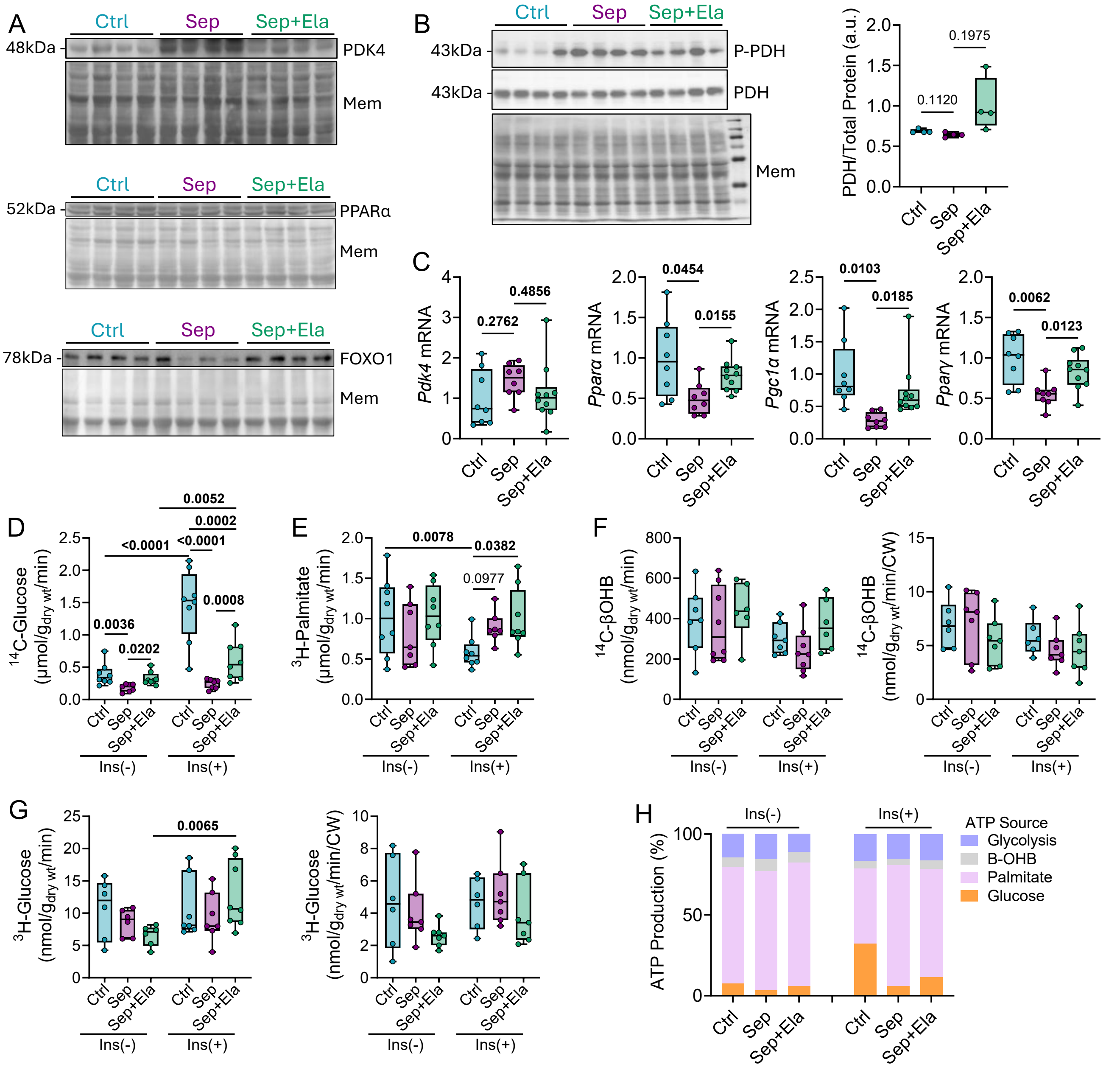

**Figure S4. Additional ex vivo working heart parameters including oxygen consumption, raw substrate oxidation rates, and proportional ATP production. A**. Representative western blots of pyruvate dehydrogenase kinase 4 levels (PDK4), peroxisome proliferator-activated receptor alpha (PPARα), and nuclear levels of FOXO1 in cardiac tissue normalized to memcode total protein stain. **B**. Representative western blots of phosphorylated PDH and total PDH normalized to memcode total protein (total PDH levels quantified on the right). **C.** Myocardial transcript levels of Pdk4, Pparα, peroxisome proliferator-activated receptor gamma coactivator 1-alpha (Pgc1α), and Ppar-γ. **D**. Glucose oxidation rates determined by tracing consumption of [U-14C]-glucose in the presence and absence of insulin not normalized to cardiac work, and **E**. Fatty acid (palmitate) oxidation rates determined by tracing consumption of [9,10-3H]-palmitate in the presence and absence of insulin not normalized to cardiac work (n=7-8). (E) **F**. βOHB (ketone body) oxidation rates measured by tracing consumption of [3-14C] βOHB, expressed as raw values (left panel) and normalized to cardiac work (right panel) (n=6-7). **G**. Glycolytic rates assessed by tracing consumption of [5-3H]-Glucose (left) and normalized to cardiac work (right). **H**. Proportional ATP production rates derived from glucose oxidation, glycolysis, palmitate oxidation, and βOHB oxidation (n=6-8). Ctrl = naive healthy control mice, Sep = cecal-slurry induced septic mice administered vehicle, Sep+Ela = cecal-slurry induced septic mice administered elamipretide. Data are presented as box-and-whisker plots showing the mean, interquartile range, minimum, and maximum values. Data were analyzed with one-way ANOVA with Sidak's multiple comparison test. Outliers were identified and excluded using the Grubbs' test with a significance level of $\alpha = 0.05$.

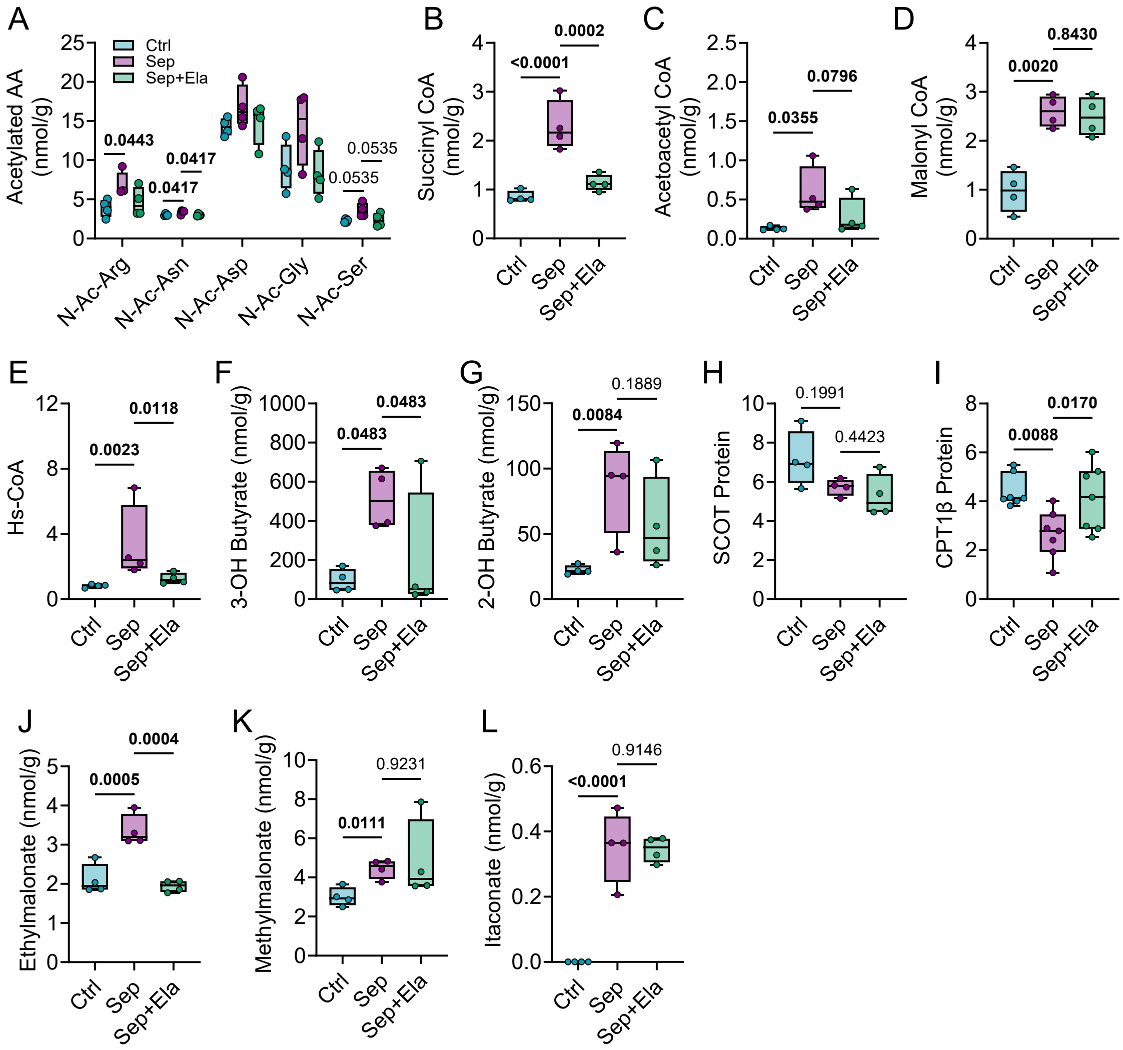


**Figure S5. A**. Myocardial abundances of acetylated amino acids (n=4). Cardiac levels of **B**. Succinyl-CoA, **C**. Acetoacetyl-CoA, **D**. Malonyl-CoA, **E**. Co-enzyme A, **F**. 3-OH Butyrate, **G**. 2-OH Butyrate. **H**. Succinyl-CoA:3-oxoacid CoA transferase (SCOT) protein and **I**. Carnitine palmitoyltransferase 1B (CPT-1β) protein levels. Myocardial levels of **J**. ethylmalonate, **K**. methylmalonate, and **L**. itaconate. Ctrl = naive healthy control mice, Sep = cecal-slurry induced septic mice administered vehicle, Sep+Ela = cecal-slurry induced septic mice administered elamipretide. n=3-4; Data are presented as box-and-whisker plots showing the mean, interquartile range, minimum, and maximum values. Data were analyzed with one-way ANOVA with Sidak's multiple comparison test. Outliers were identified and excluded using the Grubbs' test with a significance level of α = 0.05.

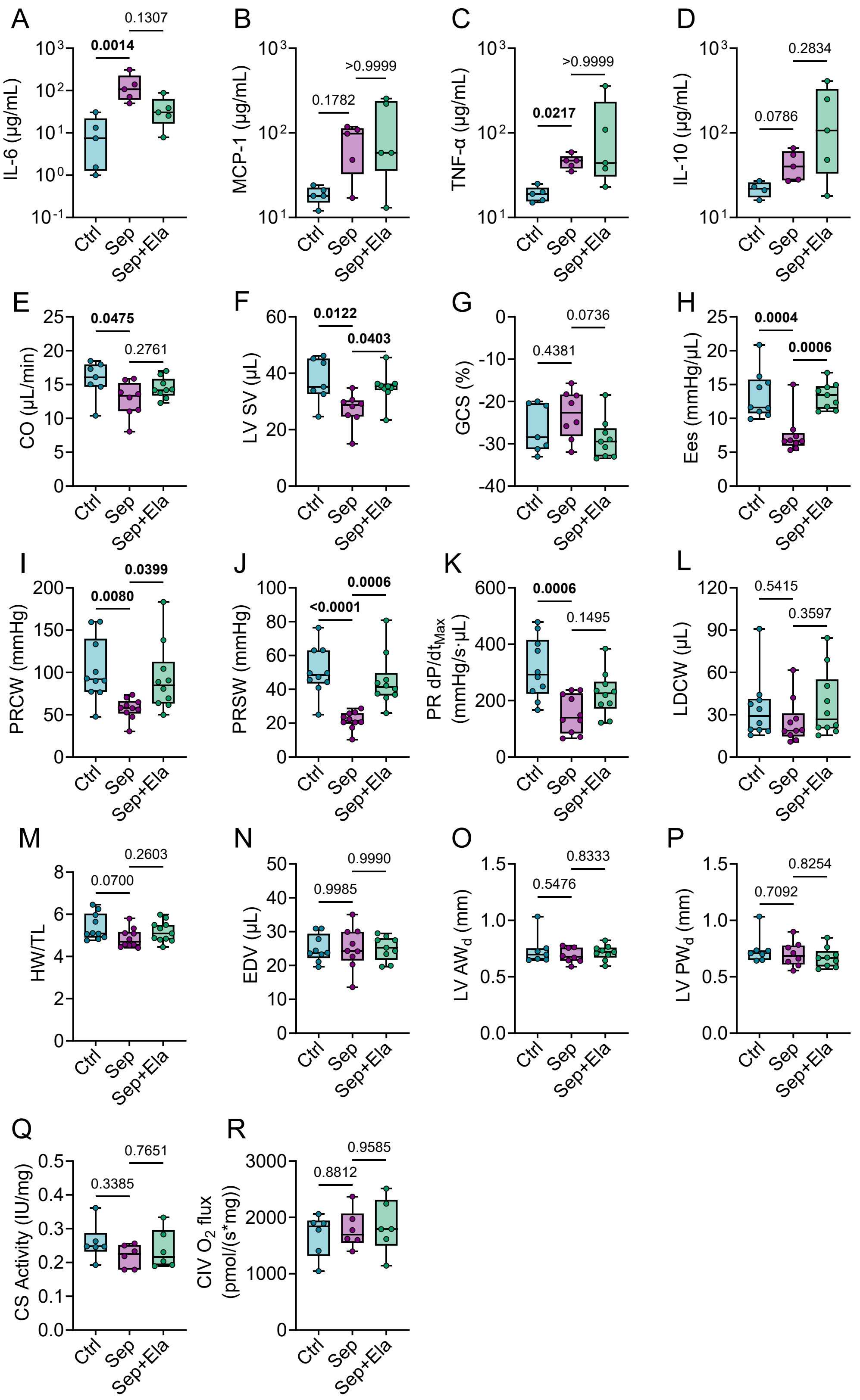


**Figure S7. Persistent cardiac and mitochondrial impairments after sepsis recovery characterized by echocardiography and pressure-volume loops. A-D**. Plasma levels of cytokines **A**. interleukin 6 (IL-

6), **B**. monocyte chemoattractant protein 1 (MCP-1), **C**. tumour necrosis factor alpha (TNF-α), and **D**. interleukin 10 (IL-10) measured10 days post-sepsis recovery. **E**. Cardiac output assessed with echocardiography 10 days post-sepsis recovery. **F**. Left ventricular stroke volume (LV SV) assessed with echocardiography (n=7-9). **G**. Global circumferential strain (GCS) assessed with echocardiography 10 days post-sepsis recovery. **H**. End systolic elastance (Ees) measured with pressure-volume loops (n=9). **I**. Preload recruitable cardiac work (PRCW) and **J**. preload recruitable stroke work (PRSW) as load-independent indicators of intrinsic contractile function assessed with pressure-volume loops. **K**. Preload recruitable (PR)dP/dTmax and **L**. Load dependent cardiac work (LDCW) determined with pressure-volume loop analyses. **M**. Heart weight (HW) normalized to tibia length (TL). **N**. End diastolic volume (EDV) measured with pressure-volume loop assessments. **O**. Left ventricular anterior wall thickness (LV AW) and **P**. posterior wall thickness (LV PW) measured during diastole with echocardiographic M-mode. **Q.** Citrate synthase activity, reflecting mitochondrial content across groups. **R**. Complex IV activity. Ctrl = naive healthy control mice, Sep = cecal-slurry induced septic mice administered vehicle, Sep+Ela = cecal-slurry induced septic mice administered elamipretide. Data are presented as box-and-whisker plots showing the mean, interquartile range, minimum, and maximum values. Data were analyzed with one-way ANOVA and Tukey's post hoc analysis. Outliers were identified and excluded using the Grubbs' test with a significance level of $\alpha = 0.05$.
